# Microtubule plus-tips act as signaling hubs for positioning the cleavage furrow during cytokinesis

**DOI:** 10.1101/339119

**Authors:** Vikash Verma, Thomas J. Maresca

## Abstract

Cell division in animal cells culminates with the formation of a contractile ring that divides the cytosol through formation of a cleavage furrow. Microtubules (MTs) are essential for furrow positioning, but the molecular nature of MT-derived spatial signals is unresolved. In this study essential cytokinesis regulators (the centralspindlin complex, aurora B kinase (ABK), and polo kinase) were visualized in *Drosophila melanogaster* (*Dm*) cells and found localize to and track MT plus-ends during cytokinesis. The RhoA GEF Pebble (*Dm* ECT2) did not robustly tip-track but became enriched at MT plus-tips rapidly following cortical contact resulting in RhoA activation and enrichment of myosin-regulatory light chain. Abrogation of cytokinesis regulator tip-tracking by EB1 depletion or deletion of a novel EB1-interaction motif (hxxPTxh) in the centralspindlin component RacGAP50C resulted in higher incidences of cytokinesis failure. We propose that EB1-dependent, MT plus-tip-based signaling hubs recruit cortical *Dm* ECT2 upon contact to locally activate RhoA.

## INTRODUCTION

Cell division ends with the formation of an actomyosin contractile ring, positioned midway between the segregated chromosomes. Constriction of the ring generates a cleavage furrow that physically divides the cytosol, a process known as cytokinesis. Establishing and maintaining the position of the cleavage furrow between daughter nuclei is vital for embryonic development, tissue and stem cell maintenance, and preserving ploidy (Li, 2007). Furthermore, cytokinesis failure yields tetraploid daughter cells and has been shown to potentiate chromosomal instability, metastatic cellular behaviors, and tumorigenesis in mice (Fujiwara et al., 2005). Therefore, a cell must ensure positioning, maintenance, and completion of the cleavage furrow with high fidelity. In animal cells, the spindle apparatus specifies the position of the cleavage furrow during anaphase (Rappaport, 1961), and it is widely accepted that the cleavage plane is positioned via microtubule (MT)-dependent mechanisms (Rappaport, 1996); however, there is lack of consensus regarding the nature and molecular composition of the MT-derived positioning signals and how they are delivered. Three prominent models, namely astral stimulation, central spindle stimulation, and polar relaxation have been proposed to contribute to defining the cleavage plane (Bringmann and Hyman, 2005; Canman et al., 2003; Cao and Wang, 1996; Crest et al., 2012; D’Avino et al., 2005; Glotzer, 2004; Hutterer et al., 2009; Mogilner and Manhart, 2016; Nguyen et al., 2014; von Dassow, 2009). The astral stimulation model proposes that positive furrow specifying signals are delivered to the cortex along equatorial astral MTs positioned in between the sister chromatids (Devore et al., 1989). Based on the assessments that equatorial astral MTs are stabilized relative to polar MTs and that MKLP1 is a plus-end directed motor, Odell and Foe (OF) developed a computational model for furrow positioning positing that equatorial astral MTs become stabilized at anaphase onset (Foe and von Dassow, 2008) and, as a result, persist long enough to serve as tracks for the motor protein MKLP1 to deliver a positive furrowing signal to the equatorial cortex (Odell and Foe, 2008). However, this assumption is not universal, and some reports indicate that contact between the cortex and the astral MTs is important for furrow initiation, and not the stabilization or dynamic state of MTs (Strickland et al., 2005a; von Dassow, 2009). In addition, there is evidence that MKLP1 motor activity is dispensable for cleavage furrow initiation (Matuliene and Kuriyama, 2002; Minestrini et al., 2003). Thus, neither the stabilization of equatorial MTs nor MKLP1 motility are universally accepted features of the astral stimulation model.

According to the central spindle stimulation model, a diffusive positive signal originating from the central spindle or midzone (an overlapping MT zone in the middle of the cell) specifies the cleavage plane. Many signaling proteins, including the chromosome passenger complex (CPC), which is comprised of four proteins: survivin, borealin, INCENP, and ABK (Carmena et al., 2012), localize to the central spindle during anaphase (Cooke et al., 1987). After anaphase onset, CPC has been shown to produce an ABK activity gradient emanating from the midzone MTs (Afonso et al., 2014; Fuller et al., 2008; Kelly et al., 2007; Sampath et al., 2004), that is generally accepted to provide a critical spatio-temporal clue for furrow positioning. An accompanying proposition for this model is that ABK facilitates the formation of centralspindlin oligomers at the equatorial plasma membrane to initiate cleavage furrow formation (Basant et al., 2015).

In favor of the polar relaxation model, it has been proposed that MT-derived inhibitory signals impede contractility near the polar cortex by blocking RhoA activity (Murthy and Wadsworth, 2008). In *C. elegans*, the nature of this inhibitory signal has recently been shown to be dependent upon TPXL-1 mediated activation of aurora A kinase (Mangal et al., 2018). Another study in *Drosophila* and human cells found that inhibitory signals originate from a kinetochore-derived PP1-Sds22 phosphatase activity gradient during anaphase (Kunda et al., 2012; Rodrigues et al., 2015).

In addition to MTs, spatio-temporal regulation of cytokinesis is controlled by a complex interplay of cyclin-dependent kinase 1 (CDK1), ABK, and polo kinase. High CDK1 activity during metaphase results in phosphorylation of the centralspindlin component MKLP1, which prevents its association with the MTs (Glotzer, 2009; Mishima et al., 2004). Once the checkpoint is satisfied and cyclin B levels drop, reduced CDK1 activity results in dephosphorylation of MKLP1, thereby allowing it to bind to MTs where it becomes highly enriched in the midzone. MKLP1 has also been shown to localize at the MT plus-ends and the cell cortex (Nishimura and Yonemura, 2006; Vale et al., 2009). However, the functional significance of MKLP1 localization to MT plus-ends is not clear. Binding and enrichment of MKLP1 on the central spindle is further regulated by ABK, which stabilizes the midzone localization of the centralspindlin complex via phosphorylation of MKLP1 (Douglas et al., 2010). In many cell types, midzone localization of the CPC is dependent on the plus-end directed motor MKLP2 (Cesario et al., 2006; Gruneberg et al., 2004; Kitagawa et al., 2013; Nguyen et al., 2014), while its cortical localization depends on actin binding by INCENP (Landino et al., 2017; Landino and Ohi, 2016). Although, the role of ABK at the central spindle is well documented; its function at the equatorial cortex is less clear and, like MKLP1, it has also been reported to localize to MT plus-tips following anaphase onset (Vale et al., 2009).

Polo kinase plays a central role in cytokinesis, where it localizes to the midzone and has also been shown to regulate many functions during mitosis (Brennan et al., 2007; Burkard et al., 2009; Carmena et al., 2014; Llamazares et al., 1991; Petronczki et al., 2008). Polo kinase phosphorylates the centralspindlin component MgcRacGAP (Ebrahimi et al., 2010), to recruit the RhoGEF, ECT2, to the midzone (Burkard et al., 2009; Petronczki et al., 2007; Somers and Saint, 2003). ECT2 produces RhoA-GTP at the membrane, which promotes cortical contractility via activation of downstream actin and myosin regulatory pathways (Bement et al., 2005; Jordan and Canman, 2012; Yuce et al., 2005). Interestingly, it was recently reported that cytokinesis can occur normally in the absence of midzone-localized ECT2 through mechanisms that signal to plasma membrane-associated ECT2 (Kotynkova et al., 2016).

PRC1 (Feo in *Drosophila*) and Kinesin-4 (Klp3A in *Drosophila*) are necessary for Polo recruitment to the central spindle in *Drosophila* and mammalian cells (D’Avino et al., 2007; Neef et al., 2007), suggesting that this mode of Polo recruitment to the central spindle might be evolutionary conserved. However, depletion of Feo does not result in cleavage furrow initiation or ingression defects (D’Avino et al., 2007), indicating that Polo recruitment to the central spindle is not necessary for furrow positioning. Nevertheless, global Polo kinase activity is essential for cytokinesis as it is required for cleavage furrow initiation (Brennan et al., 2007; Lenart et al., 2007; Petronczki et al., 2007). Taking together, these data suggest that Polo kinase and ECT2 localization to the midzone may be dispensable for furrow initiation.

In this study, we examined MT-based signaling to the process of cleavage furrow positioning in *Drosophila melanogaster* (*Dm*) S2 cells. We applied live-cell TIRF microscopy to quantitatively characterize the spatio-temporal dynamics of lynchpin cytokinesis regulators: *Dm* MKLP1 (Pavarotti in *Dm*), MgcRacGAP (RacGAP50C/Tumbleweed in *Dm*), ABK, Polo kinase, *Dm* ECT2 (Pebble in *Dm*), and active RhoA. Live-cell imaging revealed that *Dm* MKLP1, RacGAP50C, ABK, and Polo each localize to and track astral MT plus-tips within minutes of anaphase onset before becoming patterned onto a subset of equatorial astral MTs. We observed rapid recruitment and subsequent amplification of cortical Rho-GEF Pebble (*Dm* ECT2) at sites contacted by MT plus-tips, which then resulted in localized activation of RhoA at the cortex, and accumulation of myosin-regulatory light chain (MRLC). Therefore, these specialized MT plus-tips were deemed ‘’<u>c</u>ytokinesis <u>s</u>ignaling TIPs’’ (referred to hereafter as CS-TIPs). CS-TIP assembly and signaling required global, but not midzone-localized ABK activity while Polo kinase activity was not required for CS-TIP assembly, but was necessary for CS-TIPs to recruit ECT2 and activate RhoA. Cortical contact by CS-TIPs was required to maintain signaling to the contractile machinery as localized RhoA activation, and MRLC enrichment were lost concomitant with disassembly of CS-TIP components. Interestingly, localization of *Dm* MKLP1 and Polo kinase as well as efficient furrow initiation and high-fidelity cytokinesis required the plus-tip tracking protein EB1. Plus-tip tracking by the centralspindlin component RacGAP50C required a newly identified EB1-interaction motif (hxxPTxh), the deletion of which resulted in increased rates of cytokinesis failure comparable to EB1 depletion. We propose that CS-TIPs act as signaling hubs that deliver furrow positioning signals to the cortex via EB1-dependent MT plus-tip localization and tracking of centralspindlin, ABK, and Polo kinase. Cortical contact by CS-TIPs recruits *Dm* ECT2, thereby activating RhoA to aid in positioning the cleavage furrow during cytokinesis.

## RESULTS

### The centralspindlin complex, ABK, and Polo kinase localize to astral MT plus-tips following anaphase onset and become patterned onto equatorial astral MTs over time

It has long been known that the centralspindlin complex and CPC are highly enriched in the midzone during cytokinesis (Glotzer, 2009); however, previous studies in *Drosophila* and mammalian cells as well as *Xenopus* embryos have reported MT plus-tip localization of MKLP1 and MgcRacGAP (centralspindlin complex) as well as the CPC component ABK (Breznau et al., 2017; Nishimura and Yonemura, 2006; Vale et al., 2009). While there has been significant investigation of midzone populations of cytokinesis regulators, very little attention has been given to MT plus-end localized components. Thus, we sought to further investigate the spatio-temporal dynamics of the tip-localization properties of centralspindlin and ABK by live-cell TIRF microscopy of dividing *Drosophila* S2 cells expressing fluorescently tagged Pavarotti (*Dm* MKLP1), RacGAP50C (*Dm* MgcRacGAP), and ABK.

We began by imaging MKLP1, a motor protein that belongs to Kinesin-6 family (also known as Pavarotti in *Drosophila*). In metaphase, *Dm* MKLP1 diffusely localized throughout the cytosol without evident spindle MT localization **(Supplementary Movie 1)**; however, within 2-3 minutes of anaphase onset, *Dm* MKLP1 localized to and tip-tracked on the plus-ends of astral MTs (CS-TIPs) **(Figure 1A; Supplementary Figure 1E; Supplementary Movie 1)**. In agreement with prior observations (Vale et al., 2009), *Dm* MKLP1 localized uniformly to astral MT plus-tips, both polar and equatorial, before being lost from the polar tips and becoming preferentially patterned onto equatorial tips ∼10 minutes following anaphase onset **(Figure 1A; Supplementary Movie 1)**. We next visualized FP-tagged RacGAP50C since homodimers of MKLP1 and MgcRacGAP interact to form a functional centralspindlin complex (Mishima et al., 2002; Pavicic-Kaltenbrunner et al., 2007). In agreement with a recent report of MgcRacGAP localization to the MT plus-ends in *Xenopus laevis* (Breznau et al., 2017), we also observed MT plus-end localization and tip-tracking activity of RacGAP50C during cytokinesis (Figure 1B; Supplementary Movie 2). Like *Dm* MKLP1, RacGAP50C uniformly decorated polar and equatorial astral MT plus-ends within ∼2 minutes of anaphase onset before becoming patterned onto equatorial MTs **(Figure 1B; Supplementary Movie 2)**. We also observed both RacGAP50C and *Dm* MKLP1 accumulation on midzone MTs and the equatorial cortex **(Figure 1, A and B; Supplementary Movies 1 and 2)**. Thus, the two components of the centralspindlin complex, *Dm* MKLP1 and RacGAP50C, exhibit the same dynamic localization pattern on midzone MTs, the cortex, and astral MT plus-tips as cells progressed through anaphase and cytokinesis. Consistent with previous reports (Landino et al., 2017; Vale et al., 2009), we also observed ABK localization on the polar and equatorial CS-TIPs, midzone, and the equatorial cortex **(Figure 1C)**. Interestingly, ABK localization on the CS-TIPs was transient (∼1 minute) in comparison to centralspindlin components, which typically lasted for 2-10 minutes on polar MTs and >10 minutes on equatorial MTs.

**Figure 1.**
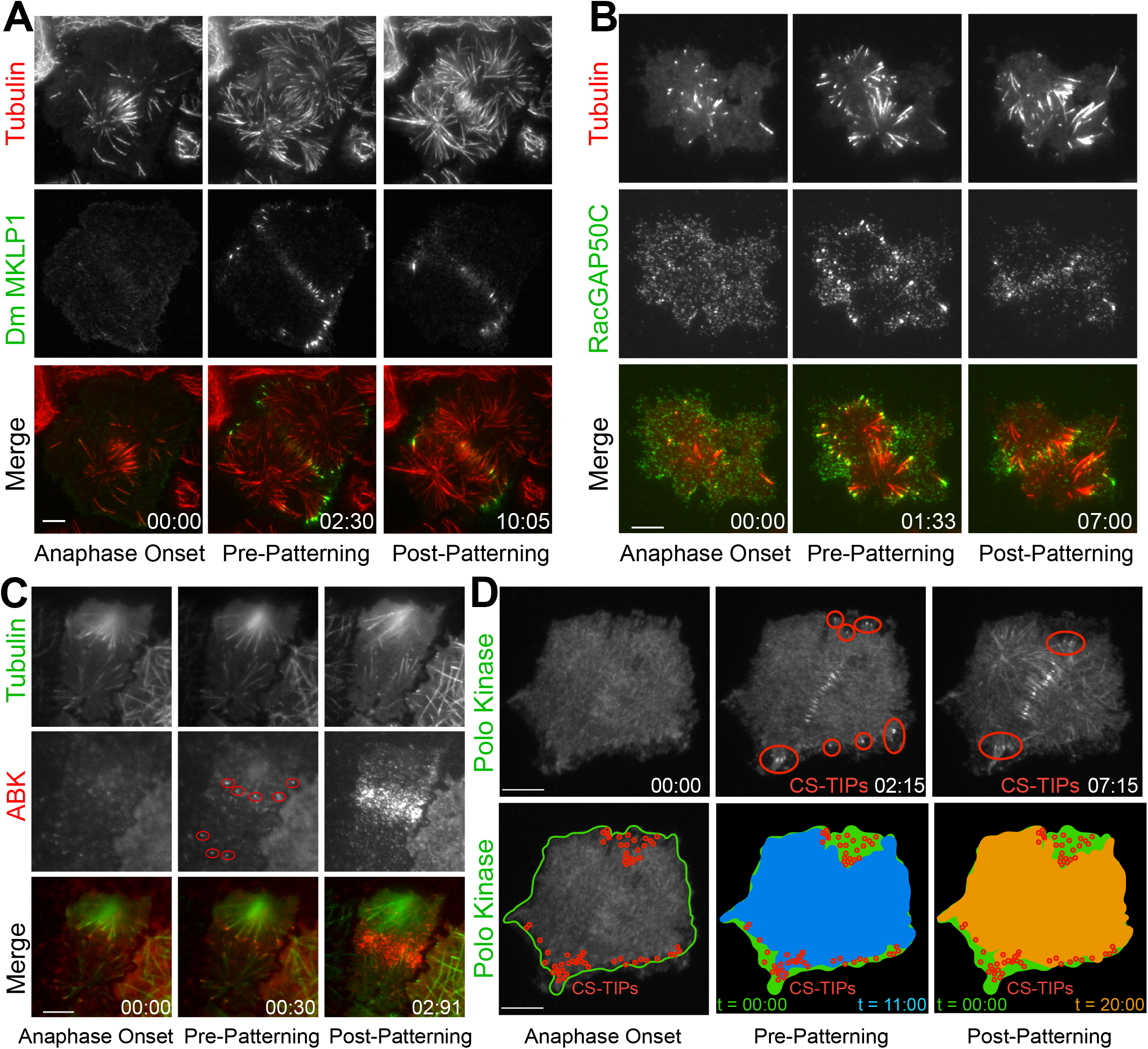
*Dm* MKLP1, RacGAP50C, ABK, and polo kinase localize to CS-TIPs and the midzone. (A) Selected still frames from the live-cell TIRF movies showing *Dm* MKLP1-EGFP (green) localization to the plus-tips of polar and equatorial MTs (red) within 2.30 minutes of anaphase onset. *Dm*-MKLP1-EGFP gets patterned on the equatorial MTs after ∼10 minutes (the last panel). (B) Selected still frames from the live-cell TIRF movies showing RacGAP50C-EGFP (green) localization to the plus-tips of polar and equatorial MTs (red) within 1.33 minutes of anaphase onset. The last panel shows patterning of RacGAP50C-EGFP mainly on the equatorial MTs. (C) ABK (red) localizes to the plus-tips of polar and equatorial MTs (green) within 30 seconds of anaphase onset. The last panel shows ABK localization to the equatorial and midzone MTs. (D) Polo localization on the MTs (red regions) within minutes of anaphase onset and then patterning on the equatorial MT tips. Outline of the cell cortex at anaphase onset with the position of CS-TIPs that appeared over the time-lapse marked with red circles (bottom panel). The membrane invaginates where CS-TIPs contact the cortex as evident in the overlays of the cell peripheries at later time-points (blue - 11:00; and orange - 20:00) relative to the cell boundary at anaphase onset (green). Zero time-points indicate anaphase onset, the next time-points indicate decoration of proteins on the polar and equatorial MTs (pre-patterning), and the last time-points indicate localization of cytokinesis proteins mainly on the equatorial MTs (patterning) in all the figures. Time: mins:secs. Scale bars, 10 μm.

Polo kinase becomes enriched on midzones during cytokinesis and its activity is required for furrow initiation and successful cytokinesis (Brennan et al., 2007; Burkard et al., 2009; Carmena et al., 1998; Lenart et al., 2007; Petronczki et al., 2007). We next applied TIRF microscopy to Polo-EGFP expressing cells to examine its spatio-temporal dynamics during cytokinesis. As expected, Polo localized to the overlapping zone of the midzone MT array **(Figure 1D; Supplementary Movie 3)**. However, Polo kinase also localized to the CS-TIPs **(Figure 1D; Supplementary Movie 3)** within ∼2-3 minutes of anaphase onset before becoming preferentially patterned onto equatorial astral tips ∼10 minutes post anaphase onset. Polo kinase uniformly localized along the length of MTs as cells advanced into late telophase, which was not surprising since Polo kinase associates with interphase MTs in *Drosophila* cells (D’Avino et al., 2007). Thus, prior to telophase, Polo kinase exhibited an identical localization pattern as the centralspindlin complex. In cells expressing Polo-EGFP we often observed inward moving zones of Polo-EGFP exclusion **(Figure 1D; Supplementary Movie 3)**. Since MTs could not penetrate these exclusion zones and were pushed inward as the zones expanded, we interpreted the Polo-EGFP exclusion zones as indicative of local regions of actomyosin contractility. Cortical contact by Polo-positive MT plus-tips consistently triggered local contractility at both the equatorial region and in the vicinity of polar astral CS-TIPs prior to patterning **(Figure 1D; Supplementary Movie 3)** suggesting that CS-TIPs may spatially trigger contractility upon cortical contact.

### CS-TIPs recruit cortical *Dm* ECT2 (Pebble) upon contact and activate RhoA

The observation of Polo-EGFP exclusion zones led us to postulate that CS-TIPs induced cortical contractility upon contact. Since cortical contractility is triggered by the GTP-bound state of the small GTPase RhoA (Bement, 2000; Glotzer, 2004; Jordan and Canman, 2012), we next examined the behavior of the RhoA-GEF ECT2 (Pebble in *Drosophila*) by visualizing dividing cells co-expressing *Dm* ECT2-EGFP and Tag-RFP-α-tubulin by TIRF microscopy. During interphase *Dm* ECT2 localized to the nucleus, but a cortical pool was also evident that remained throughout mitosis. Following anaphase onset, *Dm* ECT2 localized to midzone MTs and became enriched on the nearby equatorial cortex **(Figure 2A; Supplementary Movie 4)**. Unlike the CS-TIP components, *Dm* ECT2 did not robustly tip-track on astral MTs, rather, cortical *Dm* ECT2 co-localized with CS-TIPs within 10.5 +/− 1.0 seconds of contact **(Figure 2B)**, which was followed by amplification of *Dm* ECT2 recruitment that peaked 1.5 ± 0.1 minutes after the initial cortical contact by CS-TIPs **(Figure 2C; Supplementary Movie 4)**.

**Figure 2.**
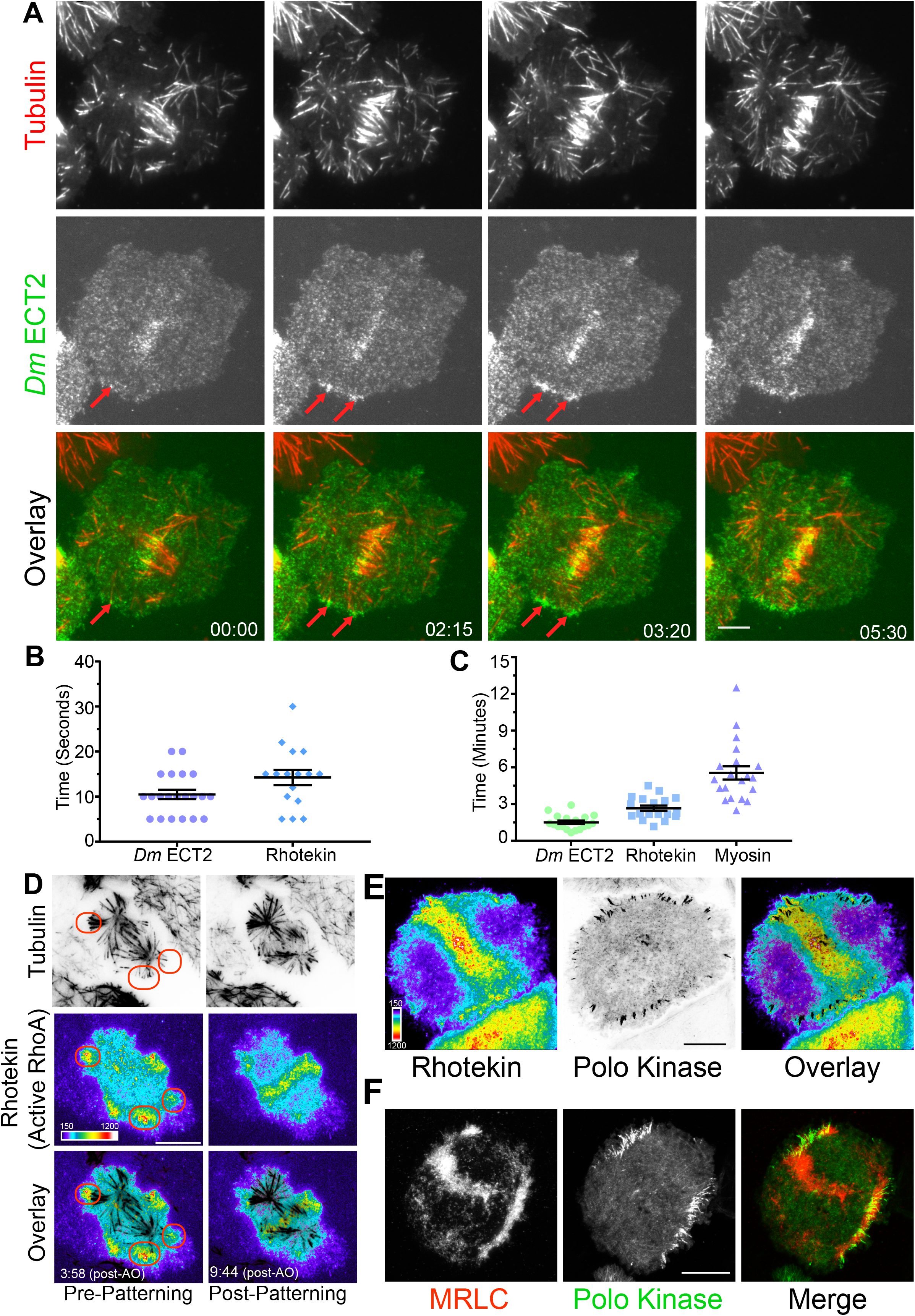
CS-TIPs, upon cortical contact recruit *Dm* ECT2 (Pebble), which then leads to RhoA activation and myosin accumulation. (A) Representative TIRF micrographs showing *Dm* ECT2 (green) localization to the CS-TIPs (red). *Dm* ECT2 localization to the CS-TIPs (time-point 00.00) is followed by its auto-amplification (later time-points) near the equatorial cortex. Red arrow heads indicate *Dm* ECT2 localization to the CS-TIPs and its amplification. (B) Dot plots showing first appearance of *Dm* ECT2 (10.48 ± 1.03), n=21 and Rhotekin (14.25 ± 1.68), n=16 after MTs contacting the cortex. Error bars: SEM. (C) Dot plots showing average peak accumulation timings of *Dm* ECT2 (1.5 ± 0.14), n=18; RhoA (2.5 ± 0.2), n=18; and Myosin (5.5 ± 0.54), n=19, Error Bars: SEM. (D) The active RhoA reporter Rhotekin is enriched near CS-TIPs within minutes (3:58) of anaphase onset (AO) and tip-signaling can precede detectable midzone RhoA activation (9:44). Red ovals denote polar signaling TIPs prior to patterning onto equatorial MTs. (E) Polo-positive CS-TIPs activate cortical RhoA (Rhotekin) upon contact during C-Phase. (F) Polo-positive CS-TIPs trigger localized cortical contractility as evidenced by MRLC accumulation. Time: mins:secs. Color wedge displays pixel values from 150-1200. Scale bars, 10 μm (B, C, and D), 5μm (A).

We next hypothesized that CS-TIP-mediated recruitment of *Dm* ECT2 should result in local activation of RhoA. TIRF-based visualization of cells co-expressing TagRFP-T-α-tubulin and Rhotekin, a reporter for active RhoA-GTP (Bement et al., 2005; Benink and Bement, 2005), N-terminally tagged with EGFP clearly revealed that a subset of astral MT plus-tips locally activated RhoA within seconds of contacting the cortex (**Figure 2, B and D; Supplementary Movie 5**). The Rhotekin (active RhoA) signal initially appeared within 14.3 +/− 1.7 seconds of cortical contact by the astral MT plus-tips and local enrichment peaked within 2.5 ± 0.2 minutes of first contact **(Figure 2C).** Due to the high spatial and temporal resolution of the live-cell TIRF microscopy, both MT plus-tip-based and midzone-derived RhoA activation signals could be visualized in Rhotekin-expressing cells. The two activation signals often arrived simultaneously, although MT plus-tip signaling sometimes preceded evident midzone-localized RhoA activation **(Figure 2D, Supplementary Movie 5)**. We further confirmed co-localization of Rhotekin (active RhoA) with Polo-positive CS-TIPs **(Figure 2E; Supplementary Movie 6)** and *Dm* MKLP1, demonstrating that CS-TIPs were responsible for localized RhoA activation.

During cytokinesis, activation of RhoA results in assembly and contraction of an actomyosin ring. We reasoned that if indeed CS-TIPs were involved in RhoA activation near the cortex, then it would result in myosin accumulation in their vicinity. To test this hypothesis, we made a stable cell line co-expressing Polo-EGFP (an indicator of CS-TIPs) and myosin regulatory light chain (MRLC) tagged with TagRFP-T-α-tubulin. Dual-color TIRF imaging of this cell line revealed a strong enrichment of MRLC near the Polo-positive CS-TIPs **(Figure 2F; Supplementary Movie 7)** that peaked within 5.5 ± 0.54 minutes of CS-TIPs contacting the cortex **(Figure 2C).** MRLC enrichment also mediated localized cortical contractility as the CS-TIPs were clearly pulled inward when the cortical contacts persisted over minutes leading to coalescence of the CS-TIPs with a band of enriched MRLC localized several microns interior to the initial CS-TIP contact points at the cortex (**Figure 2F; Supplementary Movie 7**). CS-TIP signaling was required to maintain RhoA activation and cortical contractility as polar sites of Rhotekin and MRLC enrichment were lost upon equatorial patterning and disassembly of polar CS-TIPs **(Figure 2, E and F; Supplementary Movies 6 and 7)**. Identical results for the establishment and maintenance of RhoA activation and MRLC enrichment were attained in cells expressing *Dm* MKLP1. Altogether, these results overwhelmingly support the conclusion that CS-TIPs recruit the Rho-GEF, *Dm* ECT2, to activate cortical RhoA, which results in localized myosin accumulation and induction of cortical contractility.

### Investigating the contribution of midzone components (ABK, polo kinase, kinesin-4, MKLP2) to CS-TIP assembly

Given the central role both ABK and Polo kinase play in cytokinesis, we proceeded to investigate the contribution of ABK and Polo kinase activities to CS-TIP assembly. Wash-in experiments with the *Drosophila* specific ABK inhibitor, Binucleine 2, and the Polo kinase inhibitor BI 2536 (Eggert et al., 2004; Lenart et al., 2007; Smurnyy et al., 2010) were conducted to isolate cytokinesis-specific contributions of each kinase to CS-TIP assembly. Polo-EGFP or *Dm* MKLP1-EGFP cells were continuously imaged from metaphase/anaphase onset and Binucleine 2 was directly introduced into the imaging chamber when CS-TIPs became evident. Interestingly, wash-in of Binucleine 2 resulted in a complete loss of Polo kinase and *Dm* MKLP1 from MT plus-tips within 2.5 ± 0.43 minutes of introducing the inhibitor **(Figure 3, A and B; Supplementary Movies 8 and 9).** However, wash-in experiments with the Polo kinase inhibitor BI-2536, which we and others have shown effectively inhibits *Dm* Polo kinase (Carmena et al., 2014; Kachaner et al., 2017; Lenart et al., 2007), did not affect the CS-TIP localization of either Polo kinase or *Dm* MKLP1 (Figure 3, C and D, Supplementary Movies 10 and 11). Taken together, the data demonstrate that ABK activity is required for centralspindlin and Polo kinase to localize to CS-TIPs while Polo kinase activity is not.

**Figure 3.**
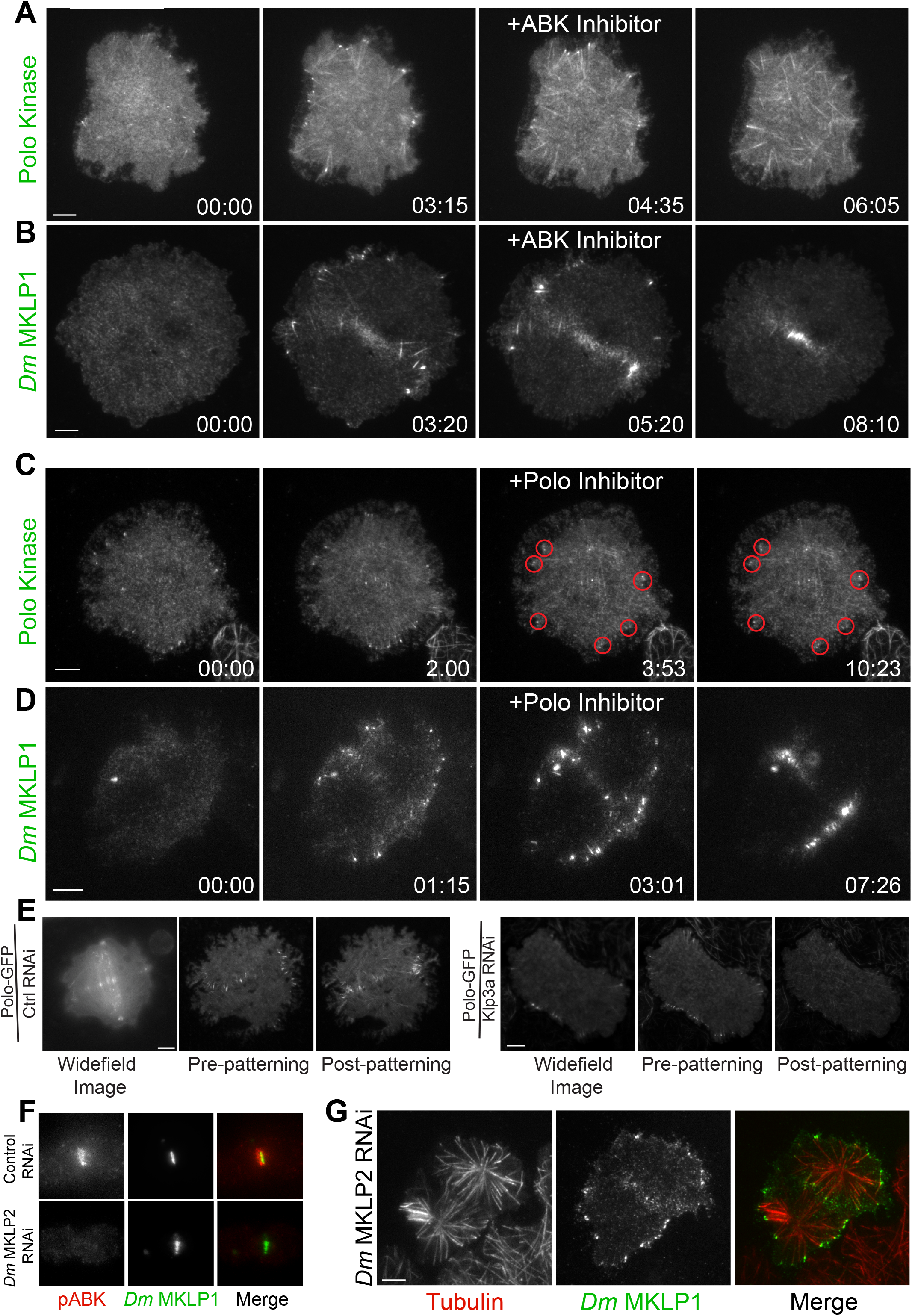
CS-TIPs require ABK activity, but not Polo kinase activity or midzone-localized CPC. (A, B) Polo and *Dm* MKLP1 are rapidly lost from CS-TIPs following wash-in of an ABK inhibitor, Binucleine 2. Zero time-points indicate anaphase onset, 04:35 and 05:20 time-points indicate the time of Binucleine 2 addition. (C, D) Polo and *Dm* MKLP1 remain localized to the CS-TIPs following wash-in of a Polo kinase inhibitor, BI 2536. Zero time-points indicate anaphase onset, 03:53 and 03:01 time-points indicate the time of BI 2536 addition. (E) Control cells show normal localization of polo-EGFP on the CS-TIPs and midzone (left). Depletion of *Dm* Kinesin-4 (Klp3A) results in loss of polo kinase from the midzone, but not from the CS-TIPs (Right). The first image is a selected frame from a wide-field movie, while last two images are selected frames from a live-cell TIRF movie for both control and Klp3A depletion. (F) Depletion of *Dm* MKLP2 leads to loss of active phosphorylated ABK (pABK) at the midzone (left panel), yet *Dm* MKLP1 assembles on CS-TIPs normally (right panel). Time: mins:secs. Scale bars, 5 μm (C,D, and E), 10 μm (A, B, and F).

PRC1 (Feo in *Drosophila*) is a well-characterized midzone MT component that preferentially bundles anti-parallel MTs (Bieling et al., 2010). PRC1 forms a complex with kinesin-4 that regulates the midzone MT length and its stabilization (Bieling et al., 2010; Hu et al., 2011; Subramanian et al., 2013; Zhu and Jiang, 2005). Interestingly, PRC1, which on its own coats individual MTs and bundles anti-parallel MTs, becomes enriched at MT plus-ends when associated with Kinesin-4 (Zhu and Jiang, 2005). *Dm* Kinesin-4 was next depleted from Polo-EGFP or MKLP1-EGFP expressing cells to investigate if the PRC1-Kinesin-4 complex contributes to CS-TIP assembly. In agreement with prior work (D’Avino et al., 2007), Polo kinase failed to localize to midzone MTs in cells with ≥ 97% of *Dm* Kinesin-4 (Klp3A in *Drosophila*) depleted **(Figure 3E; Supplementary Figure 1B; Supplementary Movie 12)**; however, Polo kinase assembled normally onto CS-TIPs **(Figure 3E).** *Dm* Kinesin-4 was next depleted from cells expressing *Dm* MKLP1-EGFP and neither midzone MT localization nor CS-TIP localization was affected **(Supplementary Figure 1A)**. These data support the conclusion that *Dm* MKLP1 and Polo kinase do not require the PRC1-Kinesin-4 complex to assemble onto CS-TIPs.

During cytokinesis, ABK localizes to midzone MTs, astral MT plus-tips, and the cortex; however, the contribution of different ABK pools to furrow formation is unclear. In many model systems, including *Drosophila*, CPC localization to the midzone is mediated by MKLP2 (Subito in *Drosophila*) (Cesario et al., 2006; Gruneberg et al., 2004; Kitagawa et al., 2013; Nguyen et al., 2014). To investigate whether midzone-localized ABK (mABK) contributed to CS-TIP assembly, *Dm* MKLP2 was depleted from cells expressing MKLP1-EGFP. Depletion of *Dm* MKLP2 resulted in a significant reduction in midzone levels of active phosphorylated ABK **(Figure 3F; Supplementary Figure 1C)**, but *Dm* MKLP1 still localized robustly to CS-TIPs and cells completed cytokinesis normally **(Figure 3G),** (Ye et al., 2015). The results from the Binucleine 2 wash-ins and *Dm* MKLP2 depletions, taken together, support the conclusion that while global ABK activity is essential for CS-TIP assembly, the mABK pool is largely dispensable **(Figure 3, A, B, F, and G)**. Since cortical ABK becomes evident following CS-TIP assembly **(Figure 1C)**, we favor the interpretation that a cytoplasmic pool of active ABK promotes assembly of CS-TIPs after anaphase onset.

### ABK and polo kinase activities are required for RhoA activation by CS-TIPs

After establishing the core components of CS-TIPs and their function in *Dm* ECT2 recruitment, and RhoA activation, we next examined the specific roles of ABK and Polo kinase in CS-TIP-mediated activation of RhoA by conducting wash-in experiments with Binucleine 2 and BI 2536 for specific inhibition of ABK and Polo kinase respectively. Dual color TIRF imaging of anaphase cells expressing TagRFP-T-Rhotekin and EGFP-α-tubulin revealed a dramatic loss of active RhoA from the vicinity of astral MT plus-tips within 3.16 +/− 0.77 minutes of introducing Binucleine 2 **(Figure 4, A and E, Supplementary Movie 13)**. Identical results were attained in cells co-expressing TagRFP-T-Rhotekin and the CS-TIP component *Dm* MKLP1-EGFP **(Figure 4C, Supplementary Movie 14)**. Loss of localized Rhotekin and *Dm* MKLP1 from MT plus-tips occurred concomitantly and within ∼ 2-3 minutes of Binucleine 2 addition, demonstrating that ABK activity is required for both CS-TIP assembly and downstream activation of RhoA.

**Figure 4.**
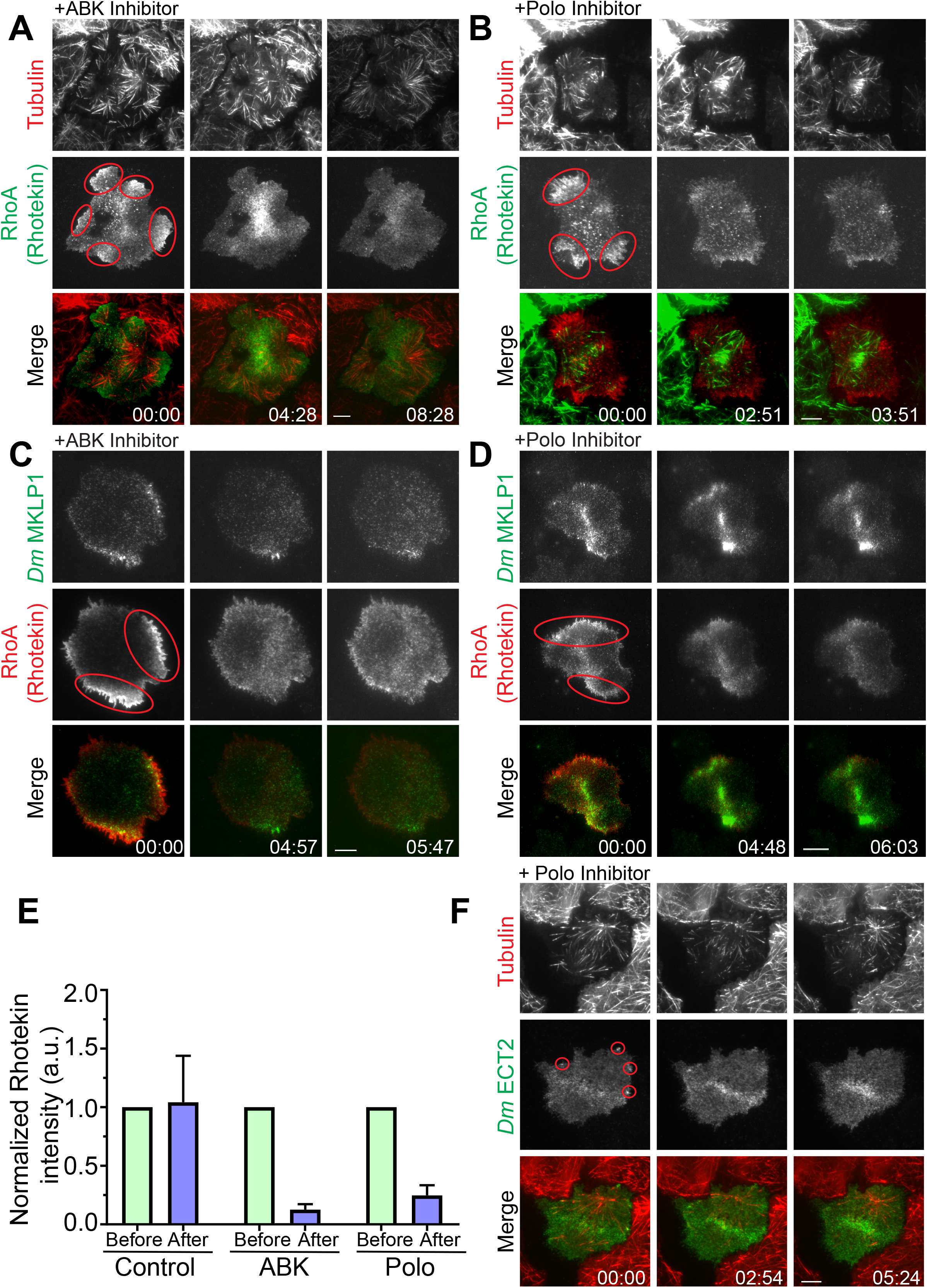
ABK and polo kinase activities are required for RhoA activation. (A, C) Active RhoA (Rhotekin) is lost from CS-TIPs following wash-in of an ABK inhibitor, Binucleine 2. (B, D) Active RhoA (Rhotekin) is lost from CS-TIPs following wash-in of Polo kinase inhibitor (BI 2536), but *Dm* MKLP1 remain on the CS-TIPs. (E) Bar graphs show normalized Rhotekin intensity before and after the addition of ABK specific inhibitor, Binucleine 2 or Polo inhibitor, BI 2536. Control cells were not treated with either Binucleine 2 or BI 2536. For quantitation of Rhotekin intensity after the addition of Binucleine 2 or BI 2536, a representative frame after ∼ 3-4 minutes was chosen. Both cortical and equatorial regions were quantified. n=4, Error bars: SD. (F) *Dm* ECT2 is lost from CS-TIPs following wash-in of Polo kinase inhibitor (BI 2536). Zero time-points indicate the time of Binucleine 2 or BI 2536 addition to the imaging chamber in all the figures. Red ovals indicate active RhoA (Rhotekin) accumulation near the CS-TIPs. Red circles indicate active *Dm* ECT2 accumulation near the CS-TIPs. Time: mins:secs. Scale bars, 5 μm.

Polo kinase activity is required for targeting ECT2 to the midzone by promoting a physical association between ECT2 and centralspindlin (Burkard et al., 2009; Petronczki et al., 2007). Thus, we next examined the effects of Polo kinase inhibition on downstream signaling to RhoA activation by CS-TIPs. Introduction of BI 2536 to anaphase cells expressing TagRFP-T-Rhotekin and EGFP-α-Tubulin resulted in a dramatic reduction in active RhoA enrichment near the astral MT plus-tips within ∼4 minutes of adding the inhibitor **(Figure 4, B and E; Supplementary Movie 15)**. The same experiment was also conducted in cells co-expressing *Dm* MKLP1-EGFP and TagRFP-T-Rhotekin to simultaneously visualize CS-TIPs and RhoA activation. In agreement with our prior observations (**Figure 2C**), Polo kinase inhibition did not affect the CS-TIP localization of *Dm* MKLP1-GFP; however, Rhotekin signal was lost in the vicinity of the *Dm* MKLP1 positive MT plus-tips **(Figure 4D; Supplementary Movie 16)**. Thus, Polo kinase activity is required for CS-TIP signaling, but not for CS-TIP assembly. We reasoned that since Polo kinase activity is required for CS-TIP-mediated activation of RhoA, then Polo inhibition should alter the localization of *Dm* ECT2 in response to physical contact by CS-TIPs. Indeed, Polo kinase inhibition resulted in the loss of localized cortical *Dm* ECT2 from the CS-TIP contact points **(Figure 4F, Supplementary Movie 17).**

### Satellite MT arrays analogous to midzone and astral MTs assemble in proximity to the cortex

Over the course of visualizing hundreds of dividing cells by TIRF microscopy we frequently noticed compelling MT structures that assembled in close proximity to the cortex as cells progressed through cytokinesis. We referred to these structures as satellite MT arrays because they formed de-novo and were not evidently linked to the spindle apparatus. Interestingly, two classes of satellite arrays were observed, which were deemed midzone and astral brush satellites because their behavior and organization were analogous to conventional midzone and astral MTs respectively.

Similar to prior observations, we noted that *Dm* MKLP1 on astral MT plus-tips bundled both parallel and anti-parallel MTs (Vale et al., 2009). Midzone satellites were comprised of anti-parallel bundles of equatorial astral MTs that crossed paths by growing towards the cortex, often buckling upon contact, and eventually becoming bundled by *Dm* MKLP1 and likely other cross-linking factors such as PRC1 **(Supplementary Movie 1).** Midzone satellites assembled in close proximity to the cortex and the ensuing cortical contractility further bundled the MTs and enriched *Dm* MKLP1 (and Polo kinase) in the overlap regions as MTs were moved inward. The process resulted in midzone satellites being in direct contact with the contractile machinery throughout cytokinesis **(Figure 5A; Supplementary Movie 1).** In many cases satellite midzones where eventually incorporated into the conventional midzone, which was typically microns away from the cortex when contractility began.

**Figure 5.**
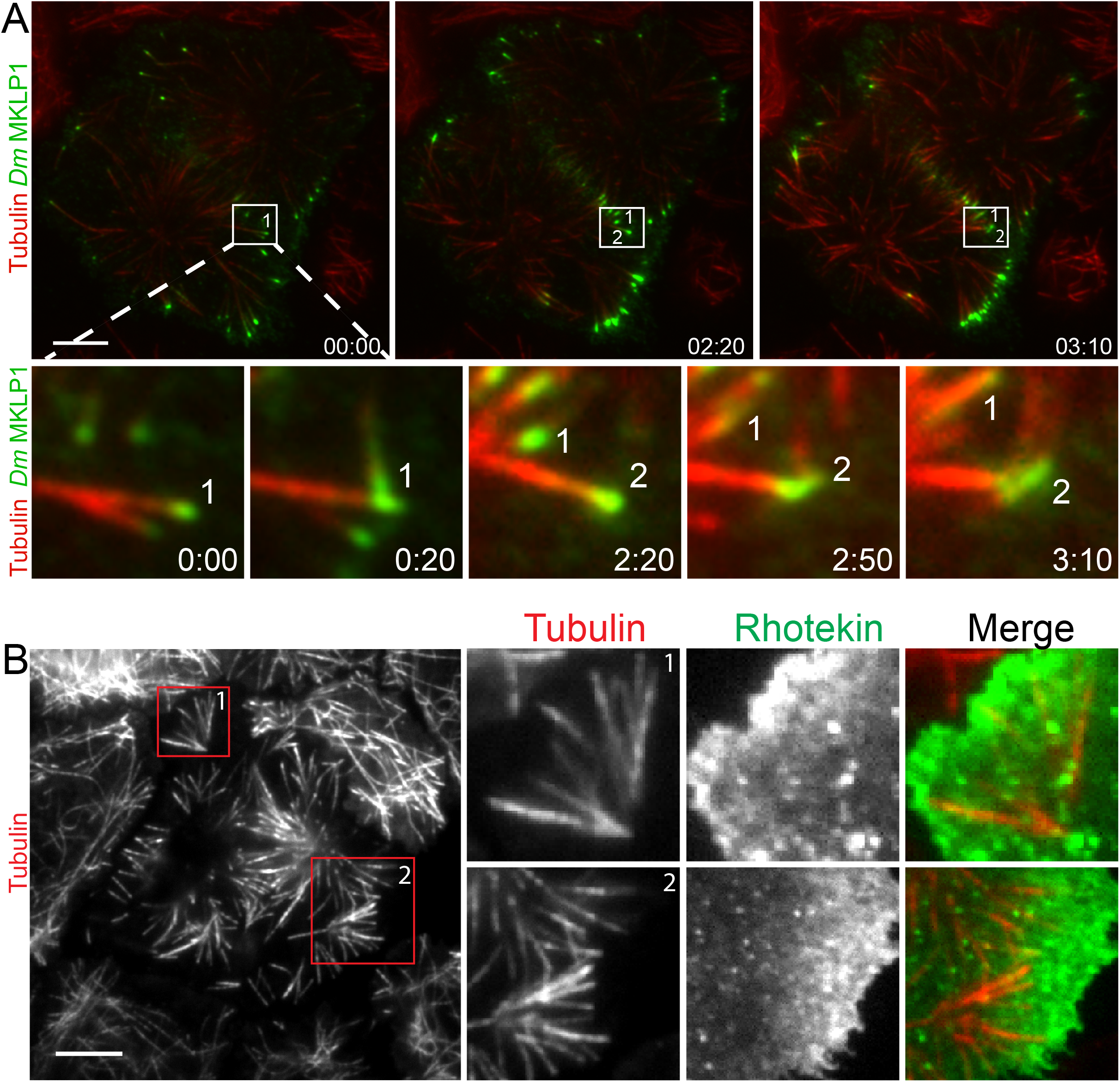
Satellite MT arrays provide a signaling platform near the cortex and midzone. (A) Representative frames from a TIRF time-lapse movie of *Dm* MKLP1-EGFP (Supplementary Video 1), (top panel). Inset shows enlarged midzone region (bottom panel). Two CS-TIP bundling events (labeled 1 and 2) occurring at the equatorial cortex. The CS-TIPs start at oblique angles, become organized into antiparallel bundles with localized *Dm* MKLP1, and are pushed inward. (B) Representative frames from a TIRF time-lapse movie of Tag-RFP-α-tubulin and EGFP-Rhotekin (left). Inset 1 and 2 show enlarged satellite MT arrays where Rhotekin gets enriched (right). Time: mins:secs. Scale bar, 5 μm.

Astral brush satellites formed de novo from cytosolic nucleation centers producing dynamic MTs that became organized over time into branched, spindle pole-like structures with MTs oriented toward the cortex **(Figure 5B).** While exhibiting astral-like properties, astral brushes typically assembled ∼5-10 microns away from the spindle apparatus and the MTs comprising astral brushes did not emanate from spindle poles or centrosome. Furthermore, the MT plus-tips of astral brushes were nearly always CS-TIPs that robustly activated RhoA, especially prior to patterning of CS-TIPs onto equatorial MTs **(Figure 5B).**

### EB1 is required for robust MT plus-tip localization of CS-TIP components and contributes to furrow initiation and successful completion of cytokinesis

Having established that CS-TIPs act as signaling hubs to activate RhoA via recruitment of cortical *Dm* ECT2, we next turned our attention to the mechanism by which CS-TIP components tip-track on astral MTs. The MT plus-end-tracking protein, EB1 is known to autonomously tip-track, and interact with other tip-tracking proteins to recruit them to polymerizing MT plus-ends (Akhmanova and Steinmetz, 2008; Duellberg et al., 2014; Honnappa et al., 2009; Strickland et al., 2005b). To test if EB1 was required for the MT plus-tip localization of CS-TIPs, EB1 was depleted by RNAi in cells co-expressing TagRFP-T-α-tubulin and either *Dm* MKPL1-EGFP; or Polo-EGFP. While >80% of control cells had robust CS-TIPs, EB1 depletions, which reproducibly yielded >90% knock down efficiencies **(Supplementary Figure 1D)**, significantly compromised tip-tracking of both Polo kinase and *Dm* MKLP1 as <10% of EB1-depleted cells exhibited MT plus-tip localization of CS-TIP components **(Figure 6, A and B; Supplementary Movie 18).** If CS-TIPs make a functional contribution to the efficiency of cytokinesis then perturbing CS-TIPs via EB1 depletion should result in cytokinesis defects. EB1 was depleted from cells expressing Tag-RFP-α-tubulin, EGFP-Rhotekin and subjected to overnight imaging. Interestingly, there was a > 4-fold increase, relative to control cells, in failure to initiate furrow ingression following anaphase onset **(Figure 6C).** Furrow initiation defects have been previously reported in dividing sea urchin eggs upon inhibitory antibody injection against EB1 or dominant negative EB1 truncation (Strickland et al., 2005b). Cytokinesis failure results in binucleated cells and, accordingly, analysis of fixed cells from EB1 depletion experiments revealed that EB1-depleted cells exhibited ∼3-fold increase in the number of binucleated cells with respect to control cells **(Figure 6D).** Thus, EB1 depletion significantly compromised CS-TIP localization and tracking, and resulted in functional consequences as cells did not initiate furrow ingression and failed cytokinesis at 3-4-fold higher rates than control cells. These results are in agreement with a prior observation in HeLa cells where depletion of EB1 and EB3 caused a 5-fold increase in cytokinesis failure (Ferreira et al., 2013)

**Figure 6.**
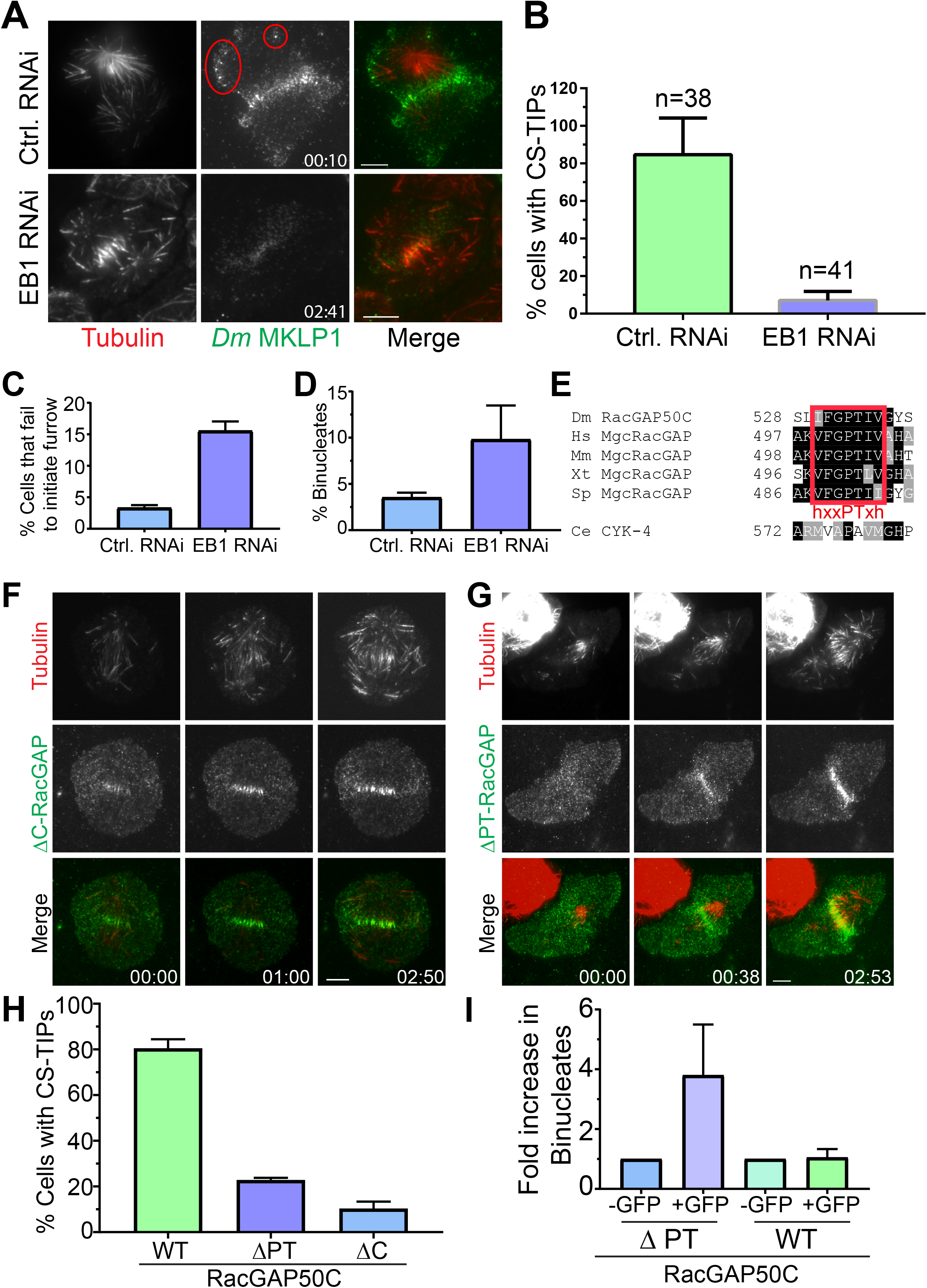
A ‘’PTIV” motif in RacGAP50C is required for localization to MT plus-tips and efficient cytokinesis. (A) TIRF micrographs showing loss of CS-TIPs assembly in cells depleted of EB1 (bottom panel); control cells show normal CS-TIPs localization (top panel). (B) Quantitation of CS-TIPs assembly in control and EB1 depleted cells (observed in both Polo and *Dm* MKLP1-EGFP-expressing cells; 2 experiments each, n = 38 and 41 for control and EB1 depleted cells respectively, Error bars: SD). (C) Quantitation of cells that fail to initiate furrowing in control and EB1 RNAi cells. n = 120 and 110 for control and EB1 RNAi cells respectively, Error bars = SD. (D) Quantitation of binucleated cells in control and EB1 RNAi, n = 410 and 452 for control and EB1 RNAi cells respectively, Error bars = SD. (E) The putative hxxPTxh motif (FGPTIV) in RacGAP50C is highly conserved although key residues in the motif are different in *C. elegans.* In the red “PT” consensus is highlighted, “h” is a hydrophobic amino acid. (F, G) Deletion of C-terminus or ‘’PTIV motif” in RacGAP50C result in loss of CS-TIPs assembly, but not midzone or cortical localization. (H) Quantitation of CS-TIPs assembly in cells expressing WT (n=21), ΔPTIV (n=26), or ΔC-RacGAP50-EGFP (n=20), 3 experiments each, Error bars: SD. (I) Bar charts showing fold increase in binucleated cells in cells expressing ΔPTIV-RagGAP50C-EGFP or WT-RacGAP50C-EGFP, n ≥ 250 for each condition, Error Bars: SD.

### MT plus-tip localization of RacGAP50C requires a new type of EB1-interaction motif that contributes to the efficiency of cytokinesis

Our characterization of the EB1-dependent nature of CS-TIP tracking is consistent with a recent work in *Xenopus laevis* embryonic cells describing the plus-tip tracking behavior of the centralspindlin component MgcRacGAP via a SxIP EB1-interaction motif during cytokinesis (Breznau et al., 2017). The SxIP motif in *X*. *laevis* MgcRacGAP is not conserved beyond *Xenopus* species despite the fact that our work and others demonstrate that centralspindlin plus-tip tracking is conserved in *Drosophila* and human cells. Our group recently identified unconventional EB1-interaction motifs in the *Drosophila* chromokinesin NOD (deemed “PT” motifs) (Ye et al., 2018, In press) that are similar to a newly characterized MT plus-tip localization sequence (Kumar et al., 2017; Manatschal et al., 2016; Stangier et al., 2018). From our work and others, we propose a PT motif consensus of hxxPTxh (“h”, hydrophobic a.a.; x, any amino acid). Sequence alignments revealed a highly conserved, possibly with the exception of *C. elegans*, putative hxxPTxh motif in the C-terminal region of MgcRacGAPs (**Figure 6E)**. After confirming RacGAP50C localization on CS-TIPs **(Figure 1B; Supplementary Figure 1F)**, we generated a C-terminal truncation of RacGAP50C lacking the hxxPTxh motif (a.a. 530-536; IFGPTIV), but retaining the ability to associate with *Dm* MKLP1 via its N-terminus (Mishima et al., 2002; Pavicic-Kaltenbrunner et al., 2007). ΔC-RacGAP50C resulted in a severe reduction in tip-tracking efficiency; however, its localization to the midzone and equatorial cortex was unaffected **(Figure 6F; Supplementary Movie 19).**

While the ΔC-RacGAP50C truncation was informative in dissecting RacGAP50’s tip-tracking activity, attributing any observed cytokinesis defects in ΔC-RacGAP50C-expressing cells to loss of CS-TIPs was complicated by the fact that ΔC-RacGAP50C prematurely truncates the GAP domain and may; therefore, compromise its GAP activity, which has been shown to be required for efficient cytokinesis (Canman et al., 2008; D’Avino et al., 2004; Minoshima et al., 2003). Therefore, we next generated a more targeted deletion mutant of RacGAP50C in which only 4 amino acids (*PTIV*^533-536^) were deleted from the hxxPTxh motif. The PTIV deletion does not remove any amino acids that are required for GAP activity and structural simulations (Biasini et al., 2014) did not reveal a significant change in the structure of the deletion mutant **(Supplementary Figure 2A).** Since attempts to make stable cell lines expressing the ΔPTIV mutant were unsuccessful, which is indicative of the mutant having a dominant negative effect, cells were examined following transient transfections. Like the ΔC-RacGAP50C mutant, ΔPTIV-RacGAP50C localized normally to midzone MTs and the equatorial cortex **(Figure 6G; Supplementary Movie 20).** However, the tip-tracking activity of RacGAP50C-ΔPTIV was compromised as the mutant exhibited tip-tracking in only ∼23% of cells that were transfected compared to ∼80% of cells expressing WT RacGAP50C **(Figure 6H**). The fact that 23% of cells expressing ΔPTIV-RacGAP50C-EGFP exhibited tip-tracking compared to <10% of EB1 depleted cells may be a consequence of homo-dimerization of the mutant with the endogenous RacGAP50C that still possesses an intact hxxPTxh motif. Nonetheless, the fact that the mutant acted in a dominant negative manner led us to closely examine the effects of its expression on cytokinesis following transient transfection.

To observe the functional consequences of PTIV deletion from RacGAP50C on cytokinesis, TagRFP-T-α-tubulin cells that were transected with ΔPTIV-RacGAP50C-EGFP were fixed ∼4 days post-transfection to quantify the number of binucleated cells. Since there was variability in the baseline levels of binucleated cells between experiments, the fold change in binucleated transfected cells (+EGFP) relative to binucleated non-transfected cells (-EGFP) cells on the same coverslip was measured for each experiment. Importantly, there was no increase in binucleated cells transfected with WT RacGAP50C-EGFP (+EGFP) compared to non-transfected cells (-EGFP) in the same population. However, transfection of ΔPTIV-RacGAP50C (+EGFP) resulted in a > 3.5-fold increase in binucleated cells compared to the baseline level of binucleated cells in non-transfected cells (-EGFP) on the same coverslip **(Figure 6I).** Taken together our data support the conclusions that furrow positioning signals travel to the cortex in an EB1-dependent manner on growing astral MT plus-ends and that this mode of delivery requires a novel and well-conserved EB1-interaction motif (hxxPTxh) in RacGAP50C that, when deleted, results in an increased incidence of cytokinesis failure.

## DISCUSSION

In this study high resolution live-cell TIRF microscopy was employed to study the molecular mechanism of furrow establishment in *Drosophila* cells. Our data demonstrate that key cytokinesis regulators (centralspindlin, ABK, and Polo kinase) localize to and track MT plus-ends within 2-3 minutes of anaphase onset. Localization of *Dm* MKLP1 and ABK has been observed previously in *Drosophila* cells but signaling capacity and regulation of these plus-end populations have not been characterized. Furthermore, the behavior of centralspindlin during cytokinesis is well-conserved as MKLP1 and MgcRacGAP localize to MT plus-tips after anaphase onset in human cells and *Xenopus* embryos (Breznau et al., 2017; Nishimura and Yonemura, 2006) thereby warranting further investigation. The application of high spatio-temporal resolution TIRF-based imaging to dividing *Drosophila* S2 cells in this study represents an ideal approach and model system for studying a conserved, but poorly characterized phenomenon.

Here we show that CS-TIP assembly is regulated in a cell cycle-specific manner. CS-TIPs assemble during a “sub-phase” of mitosis that has been referred to as cytokinesis phase (C-phase) (Canman et al., 2000). Once the checkpoint is satisfied, and CDK1 activity is reduced, centralspindlin binds to MTs, which is prevented during metaphase because of high CDK1 activity and phosphorylation of MKLP1 by CDK1 (**Figure 7, A and B**). Within minutes of anaphase onset, *Dm* MKLP1, RacGAP50C, ABK, and Polo kinase uniformly assembled on the polar and equatorial astral MT plus-tips (CS-TIPs), (**Figure 1 and 7B**). However, within ∼10 minutes CS-TIPs were lost from most of the polar astral MTs and patterned specifically on equatorial MTs (**Figure 1, 3, and 7C**). Upon physical contact, CS-TIPs recruit cortical *Dm* ECT2, which leads to local activation of RhoA and actomyosin ring assembly (**Figure 2 and 7C**). We further provide evidence that global ABK activity is absolutely required for CS-TIPs assembly, while Polo kinase activity is required for signaling. Cleavage furrow positioning signals via CS-TIPs require EB1 interaction, which is mediated by a conserved hxxPTxh motif in RacGAP50C (**Figure 6 and 7D**).

**Figure 7.**
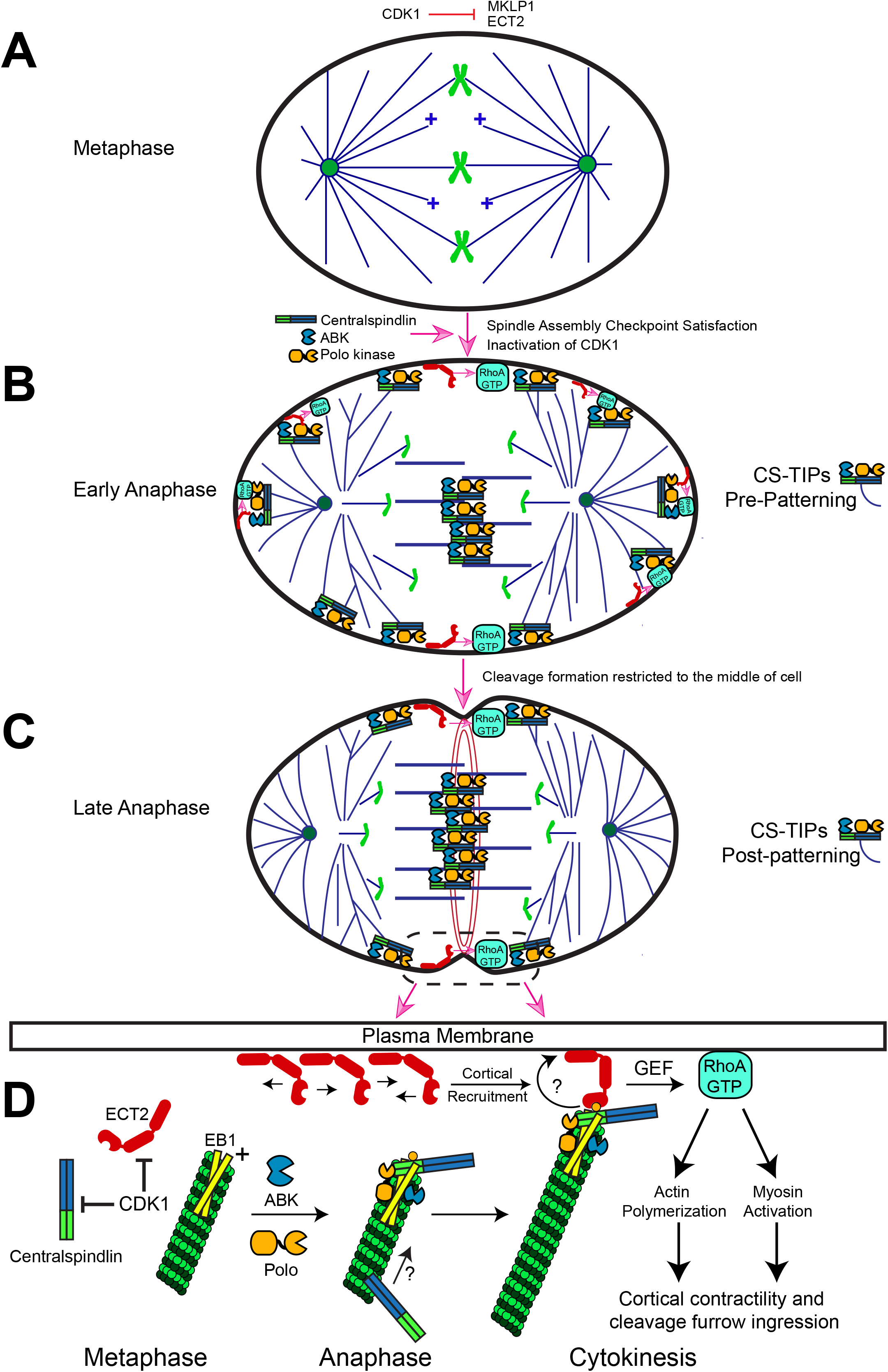
Model for the establishment of the cleavage furrow by CS-TIPs during cytokinesis. (A) Phosphorylation of MKLP1 by CDK1 during metaphase impedes its association with the spindle MTs. (B) As cell progresses towards anaphase, CDK1 activity drops, resulting in loss of MKLP1 phosphorylation and binding of MKLP1 to the spindle MTs. Within 2-3 minutes of anaphase onset, centralspindlin complex, ABK, and Polo kinase decorate the CS-TIPs. CS-TIPs after making a cortical contact, recruits Rho-GEF, ECT2, thereby activating RhoA in the vicinity. (C) CS-TIPs disassemble from the polar MT tips, and preferentially pattern on the equatorial MT tips, resulting in local activation RhoA near the equatorial cortex and assembly of actomyosin ring. (D) CS-TIPs assembly is achieved by the interaction between centralspindlin component, RacGAP50C and +TIP end-binding protein 1, (EB1). ECT2 recruitment to the CS-TIPs and its auto-amplification (shown by curved arrow and question mark) most likely involves a mechanism that requires positive feedback.

### CS-TIP- vs midzone-based signaling in positioning the cleavage furrow

MTs specify the site of furrow ingression during cytokinesis via local activation of the small GTPase RhoA. The positioning of the contractile region has been variously proposed to involve positive equatorial cues derived from the central spindle, astral MTs, or both; and inhibitory factors acting at poles (Canman et al., 2003; Mogilner and Manhart, 2016). While inferred, direct evidence of RhoA activation by astral MTs is lacking. Here we provide evidence that cortical *Dm* ECT2 and active RhoA (Rhotekin) localize to CS-TIPs within seconds of contact (**Figure 2**). *Dm* ECT2 amplification at CS-TIP cortical contact points may occur via a poorly understood positive feedback mechanism (Bement et al., 2015; Graessl et al., 2017; Tyson et al., 2003). Sustained CS-TIP signaling over a minutes time-scale further recruits myosin (MRLC) - eventually resulting in localized cortical contractility (**Figure 2D**). Due to the high spatial and temporal resolution of the TIRF-based imaging assays we employed, both MT plus-tip-proximal and midzone-derived RhoA activation signals could be readily visualized in Rhotekin-expressing cells. The CS-TIPs and midzone-derived RhoA activation signals typically occurred simultaneously, although CS-TIPs signaling sometimes preceded evident midzone RhoA activation (**Figure 2B**). The arrival of a midzone gradient-derived activation signal to the cortex would be inexorably tied to variable features of the midzone MT array such as distance from the cortex and localization/levels of any gradient-producing midzone component. To the contrary, we propose that CS-TIP-based signaling would be more consistent than midzone-mediated signaling since CS-TIPs depend primarily on MT polymerization rates, lengths, and the orientation of astral MT plus-ends, which may be derived from centrosomes or spindle poles, towards the cortex - features that are highly reproducible between anaphase spindles. In fact, the CS-TIP mode of signaling could play a more critical role in larger cells such as those in early embryonic divisions where the midzone MT array is long distances from the cortex. Although it is worth noting that in large cells amplification of the astral MT array (Mitchison et al., 2012) similar to the satellite astral brushes observed here (**Figure 5, A and B**) would also be required for astral MTs to act over very long distances.

A midzone derived ABK gradient has been proposed for cleavage furrow positioning (Fuller et al., 2008; Tan and Kapoor, 2011); and in addition to ABK, Polo kinase has also been shown to induce furrow initiation by recruiting RhoGEF, ECT2, to the central spindle (Petronczki et al., 2007), indicating that both ABK and Polo activity is required for furrow positioning. However, a complete understanding of how ABK and/or Polo kinase activate RhoA to initiate furrowing is still lacking. We show that CS-TIPs assembly and signaling was unaffected by treatments that mis-localize midzone pools of active ABK (*Dm* MKLP2 depletion, Figure 3) and Polo kinase (*Dm* Kinesin-4 depletion, Figure 3E). While global inhibition of either ABK or Polo kinase activities leads to severe cytokinesis defects, furrow positioning and cytokinesis was not evidently impacted in this study by mis-localization of midzone pools of ABK or Polo kinase (Figure 3). Thus, midzone-based ABK and polo kinase activity gradients are either dispensable for furrow positioning in *Drosophila* or redundant with CS-TIP-based furrow positioning cues and/or other pathways. Our observation further supports a recent report where artificial targeting of ECT2 was sufficient to restore cytokinesis, and mutations that prevented midzone localization of ECT2 didn’t result in any apparent cytokinesis defects in most of the cells (Kotynkova et al., 2016).

### Establishment and maintenance of cleavage furrow positioning signals

The formation of actomyosin ring and contractility typically lasts for minutes, while the furrow must be positioned correctly, it must also be maintained for successful cytokinesis. During our high-resolution imaging, we often observed ‘‘satellite MT arrays’’ that were in close proximity to the cortex and assembled after contractility had been established. Near the equatorial cortex, midzone satellites were built from equatorial astral MTs that became bundled and forced into anti-parallel MT arrays by the contractile machinery. Importantly, once contractility was established, the midzone satellites maintained direct physical contact with the cortex such that any midzone-based cues would no longer have to act over long distances to signal to the cortex. The other type of satellite MT array - astral brushes - also assembled after contractility had begun, but via apparent branching MT nucleation from astral MTs. The astral brushes, in which MT plus-ends were oriented toward the cortex, locally activated RhoA by creating a high density of new CS-TIPs. We posit that midzone and astral brush MT satellites could fulfill two critical roles during cytokinesis as autonomous signaling centers that can 1) act at distances far from their analogous structures (midzones and poles/centrosomes) in the spindle proper, and 2) maintain contractility after it has been established.

### Role of EB1 and a conserved hxxPTxh motif in RacGAP50C in cleavage furrow positioning

We found that localization of the centralspindlin complex on the CS-TIPs is dependent on EB1. This is consistent with recent work in *Xenopus laevis* embryonic cells describing the plus-tip tracking behavior of the centralspindlin component MgcRacGAP via a SxIP EB1-interaction motif during cytokinesis (Breznau et al., 2017). However, the SxIP motif in X. *laevis* MgcRacGAP is not conserved beyond *Xenopus* species despite the fact that centralspindlin plus-tip tracking is well conserved (Nishimura and Yonemura, 2006; Vale et al., 2009). We recently identified unconventional EB1-interaction motifs (“PT” motifs) in the *Drosophila* chromokinesin NOD (Ye et al., 2018, In press) that are similar to a newly characterized MT tip localization sequence (Kumar et al., 2017). Interestingly, sequence alignments revealed a highly conserved putative hxxPTxh motif in the C-terminal region of the MgcRacGAPs. Depletion of EB1, truncation of RacGAP50C C-terminus, and deletion of PTIV from the putative hxxPTxh motif from the RacGAP50C yielded the same phenotypes - loss of plus-end localization and tracking of centralspindlin in a significant majority of cells and an increase incidence of cytokinesis failure. Our results are consistent with earlier reports where injection of an inhibitory antibody against EB1 or dominant negative EB1 truncation into dividing sea urchin eggs resulted in delayed furrow ingression (Strickland et al., 2005b), and reported finding of EB1 and EB3 depletion in HeLa cells that caused a 5-fold increase in cytokinesis failure (Ferreira et al., 2013). EB1 depletion in *Drosophila* S2 cells has also been reported to cause spindle defects (Rogers et al., 2002), however we did not observe any obvious spindle morphology defects. Although, we reproducibly achieved ≥ 90% EB1 depletion, this discrepancy could result from different levels of EB1 depletion. While similar differences in the extent of mitotic phenotypes were reported in the recent knockout studies of EB proteins in HeLa cells (McKinley and Cheeseman, 2017; Yang et al., 2017), these studies didn’t focus on cytokinesis, so it will be worthwhile to investigate whether the hxxPTxh motif is required for centralspindlin plus-tip localization and efficient cytokinesis in other cell types.

### CS-TIP assembly, signaling, and patterning onto equatorial MTs

The observation that *Dm* MKLP1, RacGAP50C, ABK, and Polo localize to the plus-ends of polar and equatorial astral MT tips before becoming patterned onto equatorial astral MTs raises an important question: how do CS-TIPs selectively disassemble from the polar astral MTs? Since we have shown that CS-TIPs activate cortical contractility, the patterning mechanisms that disassemble polar CS-TIPs would be akin to a polar relaxation mechanism. A recent study in *C. elegans* found that TPXL-1 *(C. elegans* TPX-2 homologue) mediated activation of Aurora A kinase was responsible for clearing anillin and F-actin (Mangal et al., 2018) from the polar cortex. A polar Aurora A kinase activity gradient that regulates kinetochore-MT attachments has been visualized in *Drosophila* and mammalian cells (Chmatal et al., 2015; Ye et al., 2015) although its role in C-phase warrants further study. Also in *Drosophila*, a kinetochore based PP1 phosphatase activity gradient derived from a PP1-SDS22 complex was proposed for the loss of actin and inactivation of ezrin/radixin/moesin proteins at the polar cortex (Kunda et al., 2012; Rodrigues et al., 2015). It is entirely possible that phosphatase and/or kinase activity gradients in *Drosophila* contribute to the loss of CS-TIPs from polar astral MTs. Alternatively, a chromosome based RanGTP gradient has been shown to locally reduce Anillin at the cortex (Kiyomitsu and Cheeseman, 2013), and could also impede CS-TIPs-based signaling from the polar astral MTs as the chromosomes approach the polar cortex in late anaphase prior to nuclear envelope reformation. RanGTP regulates many functions during cell division, including nuclear-cytoplasmic transport and spindle assembly (Kalab et al., 2006; Kalab et al., 2002; Wu et al., 2013), and because of this, it is difficult to isolate specific temporal Ran-dependent functions during cytokinesis. A small molecule inhibitor, importazole (2,4-diaminoquinazoline) has been shown to specifically disrupt RanGTP and importin-β interaction in *Xenopus* egg extracts and cultured cells (Soderholm et al., 2011), and could potentially be used to isolate specific temporal functions during cytokinesis. However, cytokinesis can occur in the absence of chromatin (Alsop and Zhang, 2003; Rappaport, 1996). While we presently favor the hypothesis that polar- or chromosome/kinetochore-derived activity gradients regulate CS-TIP patterning, a more detailed investigation is necessary to resolve the question of how polar CS-TIPs are selectively disassembled as cells progress through cytokinesis.

## MATERIALS AND METHODS

### Cloning and site directed mutagenesis

The *Drosophila* MKLP1 gene (known as Pavarotti in *Drosophila*, CG1258) and ECT2 (kwon as Pebble in *Drosophila*, CG8114) were PCR amplified from cDNA clones RE22456 and SD01796 respectively with a 5’ Spel site and a 3’ Xbal site. The resulting PCR products of the individual genes were then inserted into the 5’ Spel and 3’ Xbal sites of the pMT/V5 His-B vector (Invitrogen) containing in-frame EGFP gene at the 3’ end, cloned between 5’ Xbal and 3’ Sacll sites, and the CENPC promoter at the 5’ end, cloned between single Kpn1 site. The regulatory light chain of the non-muscle type 2 myosin (known as Spaghetti squash in *Drosophila*, CG3595) was amplified from the cDNA clone LD14743. The resulting PCR product was cloned between 5’ Spel and 3’ Xbal of the pMT vector containing in-frame RFP gene at the 3’ end and Mis12 promoter at the 5’end. EGFP-Rhotekin was PCR amplified from the pCS2-eGFP-rGBD plasmid (gift from the Bement lab (UW-Madison) via Wadsworth lab (UMass Amherst)). The resulting PCR product was cloned between 5’ Xhol and 3’Apal sites of the pMT vector containing CENPC promoter at the 5’end cloned between single KpnI site. A second version of Rhotekin was also created in the pMT vector where we swapped the EGFP fluorophore with RFP. Polo kinase under its own promoter was cloned between 5’ Kpnl and 3’ Spel sites of the pMT vector containing in-frame EGFP gene, cloned between 5’ Spel and 3’ Apal sites. The RacGAP50C (also known as Tumbleweed in *Drosophila*, CG13345) gene was amplified from genomic DNA of *Drosophila.* The resulting PCR product was cloned in a similar fashion as described for MKLP1. A four-amino acid deletion (*PTIV*^533-536^) in the RacGAP50C gene was generated by Q5 site directed mutagenesis kit (NEB, MA, USA), following manufacturer’s protocol. C-terminal truncation of the RACGAP50C gene was created by amplifying amino acids (1-526) from the full length RacGAP50C (a.a. 1-625) gene. The resulting PCR product was inserted into the 5’ Spel and 3’ Xbal sites of the pMT vector containing in-frame EGFP gene at the 3’ end, and the CENPC promoter at the 5’ end. See Table 1 for the primers used in cloning.

**Table 1.**
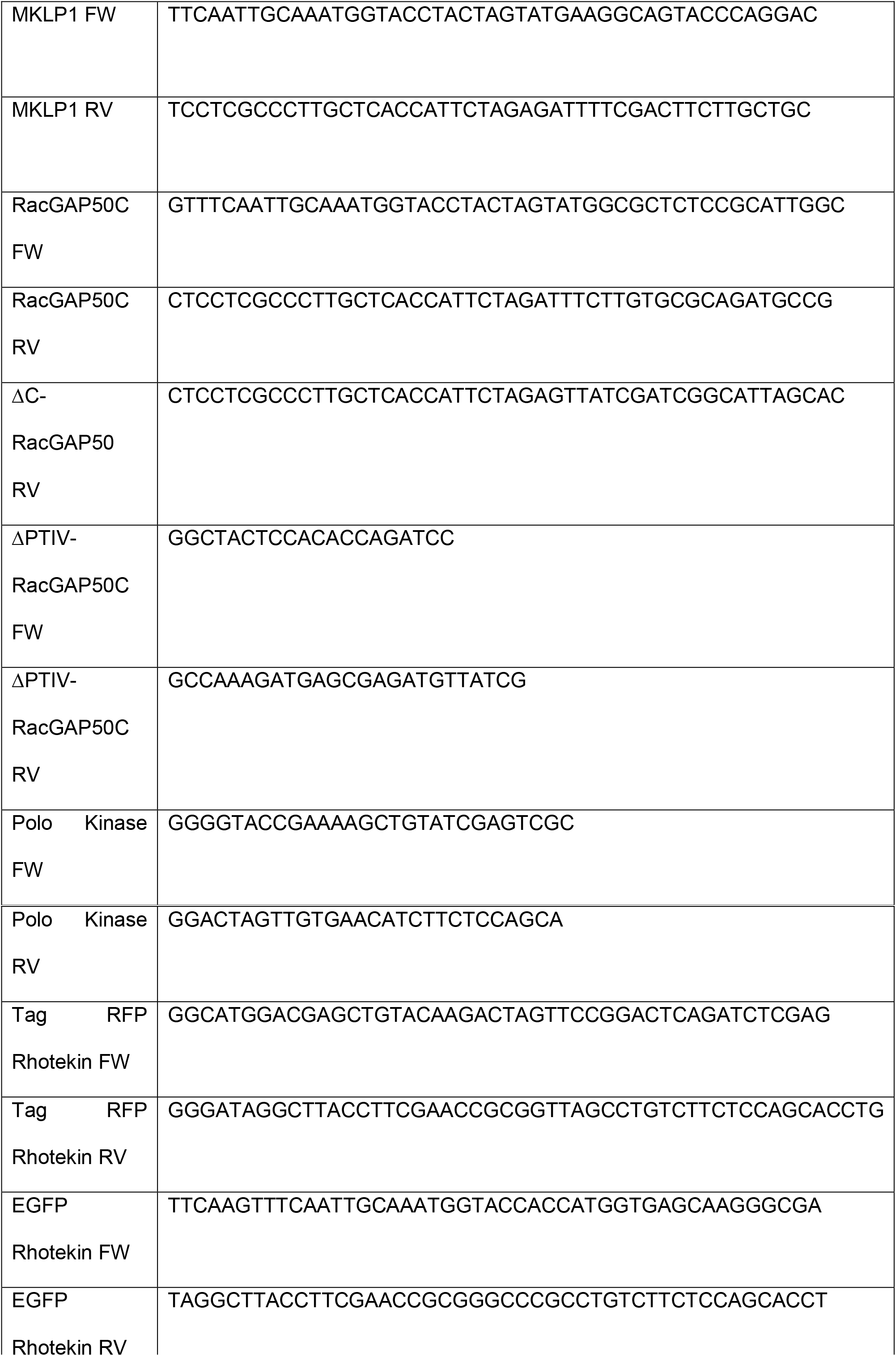
Primers for cloning

### *Drosophila Schneider (S2)* cell culture and generation of transient/stable cell line

All cell lines were grown at 25 °C in Schneider’s medium (Life Technologies, Carlsbad, CA), containing 10% heat-inactivated fetal bovine serum (FBS) and 0.5X antibiotic-antimycotic cocktail (Gibco, Life Technologies, NY, USA). Cell lines were generated by transfecting the DNA constructs containing the gene of interest with Effectene transfection reagent (Qiagen, Hilden, Germany), following the manufacturer’s protocol. After 4 days of transfection, expression of EGFP/RFP tagged proteins was checked by fluorescence microscopy. To make a stable cell line, cells were selected in the presence of Blasticidin S HCl (Thermo Fisher Scientific, Waltham, MA) and/or Hygromycin (Sigma-Aldrich) until there was no observable cell death. Thereafter, cell lines were either frozen down or maintained in the S2 media @ 25 °C without Blasticidin and/or Hygromycin B. Transient cell lines were imaged after 4-5 days of transfection, and they were not subjected to antibiotics selection. ABK-GFP and mCherry-α-tubulin cell line, was a generous gift from Dr. Eric Griffis, University of Dundee.

### RNA interference (RNAi) experiments

Around 500 base pairs of DNA templates containing T7 promoter sequence at the 5’ end for *Dm* MKLP2 (also known as Subito in *Drosophila*, CG12298), EB1 (CG3265), and *Dm* Kinesin-4 (also known as Klp3A in Drosophila, CG8590) were generated by PCR. Double-stranded RNAs (dsRNAs) were synthesized from the respective DNA templates at 37 °C using the T7 RiboMax Express Large Scale RNA Production System (Promega Corp., Madison, WI), following the manufacturer’s protocol. For RNAi experiments, cells at about 25% confluency were incubated in a 35 × 10 mm tissue culture dish for an hour. Thereafter, media was carefully aspirated off the dish and 1 ml of serum-free S2 media containing 20 *μ*g of dsRNA was added to the dish. After 1 hour, 1 ml of fresh S2 media containing FBS was added to the dish and incubated for 2-4 days at 25 °C. See Table 2 for primer sequences.

**Table 2.**
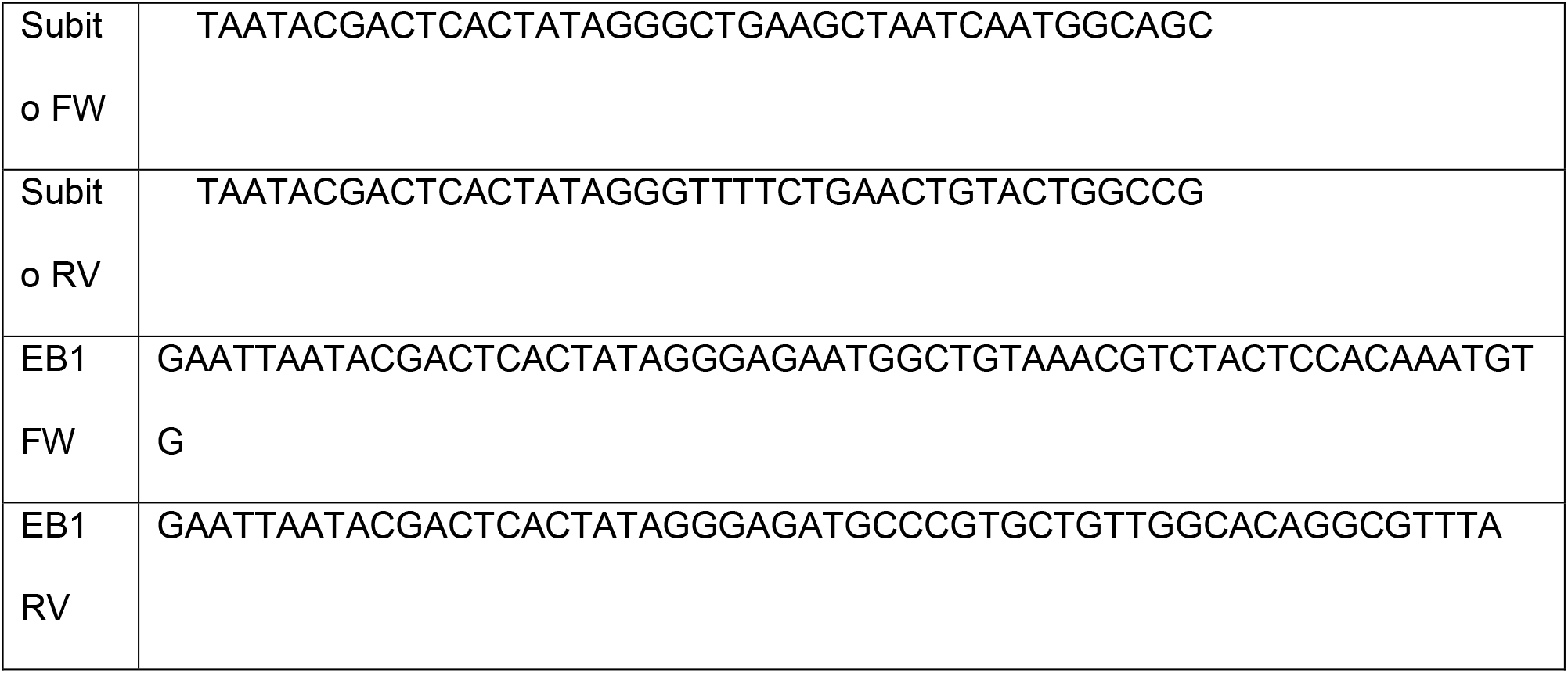
Primers for RNAi

### Immunofluorescence

*Drosophila* S2 cells were plated on an acid-washed Concanavalin A (Sigma-Aldrich) coated coverslips for 30 minutes, then cells were quickly rinsed with 1X BRB80 buffer. Subsequently, cells were fixed with 10% paraformaldehyde for 10 minutes. Cells were then permeabilized with phosphate-buffered saline (PBS) containing 1% Triton X-100 for 8 min, rinsed 3 times with PBS plus 0.1% Triton X-100, and blocked with 5% boiled donkey serum ((Jackson Immuno Research Laboratories, Inc., West Grove, PA) for 60 minutes. All primary antibodies were diluted in 5% boiled donkey serum. Anti-Phospho-aurora A/B/C (Cell Signaling Technology, Danvers, MA) was used at a concentration of 1:1000, and anti-tubulin antibody (DM1α Sigma-Aldrich) at 1:2000. All secondary antibodies (Jackson Immuno Research Laboratories, Inc., West Grove, PA) were diluted in boiled donkey serum at 1:200. After secondary antibody incubation, coverslips were washed 3 times with PBS plus 0.1% Triton X-100, followed by incubation with 4’,6’-diamidino-2-phenylindole (DAPI) at a concentration of 1:1000 for 10 minutes, and 3 additional washes with PBS plus 0.1% Triton X-100. Coverslips were sealed in a mounting media containing 20 mM Tris, pH 8.0, 0.5% N-propyl gallate, and 90% glycerol. Four-color Z-series consisting of ∼30 planes at 0.2-μm intervals were acquired for GFP, RFP, Cy5, and DAPI channels on a TIRF-Spinning Disk Confocal system assembled on a Nikon Ti-E microscope equipped with a 100X 1.4 NA DIC Apo Oil immersion objective, two Hamamatsu ORCA-Flash 4.0 LT digital CMOS camera (C11440), 4 laser lines (447, 488, 561, 641), and a MetaMorph software. Fluorescence intensities were obtained by drawing a 50 × 100 grid manually around the maximum-intensity projection of the Z-series images. The local background was estimated by placing the 50 × 100 region to a nearby place. To obtain the ratio intensities, identical regions were drawn manually on the *Dm* MKLP1-EGFP channel and transferred to a phosphorylated ABK (pABK) channel.

### Quantitative Western blotting

Equal amount of proteins was loaded into a 10 or 8% SDS-PAGE gel. After running the gel, proteins were transferred to a nitrocellulose membrane using the Trans-Blot Turbo transfer system (Bio-Rad Laboratories, Inc., Hercules, CA) for 10 minutes. Subsequently, the membrane was incubated in 5% milk (V/V made in Tris-buffered saline with 0.1% Tween) for one hour. Following blocking, the membranes was incubated with their respective primary antibodies (α-EB1 - 1:5000, α-Klp3A - 1:500, α-Tubulin - 1: 10,000) for 1 hour, followed by 3 times wash and secondary antibody incubation at 1:5000 dilution for 1 hour. Following secondary antibody incubation, membrane was washed 3 times in Tris-buffered saline with 0.1% Tween (TBST) and imaged with a G:BOX system controlled by GeneSnap software (Syngene, Cambridge, U.K.). Images were further quantified to estimate the knock down efficiency with the help of Fiji/Image J software (Schindelin et al., 2012). To obtain the intensity values, identical regions were drawn over all the bands and their integrated intensity was recorded. Intensity values were normalized to their respective loading controls (Tubulin) to estimate the knockdown efficiency.

### Fluorescence and Live-Cell Total internal reflection fluorescence (TIRF) microscopy

Around 500 to 700 μl of cells, expressing the gene of interest were seeded on a 35mm glass bottom dish with 20mm bottom well (Cellvis, CA, USA) coated with Concanavalin A for 30-60 minutes. Before imaging 2 ml of fresh S2 media containing FBS was added to the dish. Live-cell TIRF movies were acquired on a Nikon Ti-E microscope equipped with a 100X 1.4 NA DIC Apo Oil immersion objective, a Hamamatsu ORCA-Flash 4.0 LT digital CMOS camera (C11440), 4 laser lines (447, 488, 561, 641), and a MetaMorph software. Live-cell TIRF movies of all the constructs (MKLP1-EGFP and Tag-RFP-α-tubulin, RacGAP50C-EGFP and Tag-RFP-α-tubulin, Polo Kinase-EGFP, ABK-GFP and Tubulin mCherry, EGFP-Rhotekin and Tag-RFP-α-tubulin, RFP-Rhotekin and *Dm* MKLP1-EGFP, RFP-Rhotekin and MRLC-EGFP, *Dm* ECT2-EGFP and Tag-RFP-α-tubulin) were acquired at the following settings: 561 exposure time 200ms, 488 exposure time 200ms, temperature 25 °C, and frame acquisition rate 5 Sec. To observe cytokinesis defects, overnight imaging was performed on control and EB1 depleted cells. Cells were seeded on a Concanavalin A coated 35mm glass bottom dish as described above. A total of 76 time-points were acquired for 50 stage positions at the interval of 4 minutes @ 25 °C on a Nikon Eclipse Ti-E microscope (Nikon, Tokyo, Japan) equipped with 40X/1.30 Oil Nikon Plan Fluor DIC N2 objective (Nikon), Andor iXon_3_ EMCCD camera (Andor Technology, Belfast, U.K.), and Metamorph software (Molecular Devices, Sunnyvale, CA). Images were further analyzed with the help of MetaMorph software.

### Chemical perturbation experiments

Cells expressing *Dm* MKLP1-EGFP; Tag-RFP-α-Tubulin, Polo-EGFP, Rhotekin-EGFP; Tag-RFP-α-Tubulin, Rhotekin-RFP; *Dm* MKLP1-EGFP, *Dm* ECT2-EGFP; Tag-RFP-α-Tubulin were subjected to ABK or Polo inhibition. Cells were seeded on a Concanavalin A coated 35mm glass bottom dish with 20mm bottom well (Cellvis, CA, USA) for 30-60 minutes. Before imaging 2 ml of fresh S2 media containing FBS was added to the dish. Live-cell TIRF movies were acquired on a Nikon Ti-E microscope as described above. The whole field of view under the microscope was scanned to find mitotic cells. Mitotic cells expressing the gene of interest were imaged at the interval of 20-30 seconds until the anaphase onset. After anaphase onset, when the CS-TIPs were visible, S2 media in the glass bottom dish was exchanged with S2 media containing Binucleine 2 (40μM) for ABK inhibition or BI 2536 (1μM) for Polo inhibition. Thereafter, imaging was continued at 5 second intervals. Images were further analyzed with the help of Metamorph software (CA, USA).

### GFP/RFP intensity quantitation

GFP/RFP intensities before and after the addition of inhibitor (Binucleine 2 or BI 2536) were quantified to estimate the effects of inhibitor on CS-TIPs patterning and signaling. Identical regions were drawn over the GFP/RFP puncta to measure the fluorescence intensity. The local background was estimated by placing the same region to a nearby place. Background intensity values were subtracted from GFP/RFP values to estimate the actual GFP/RFP fluorescence intensity. Histograms and dot plots were generated using Prism software (Graphpad, CA, USA), and figures were assembled using Adobe Illustrator software (Adobe, CA, USA).

## ACKNOWLEDGMENTS

Authors would like to acknowledge Patricia Wadsworth, UMass and Alex Mogilner, NYU. We are also grateful to Eric Griffis, University of Dundee, for the kind gift of ABK cell line; and Bill Bement, UW-Madison for the kind gift of Rhotekin plasmid via Patricia Wadsworth. This work was supported by an NIH grant (5 R01 GM107026) to TJM.

## AUTHOR CONTRIBUTIONS

T.J.M conceived the project. T.J.M and V.V designed and performed the experiments, analyzed the data, and wrote the paper.

## SUPPLEMENTARY FIGURES

**Supplementary Figure 1.**
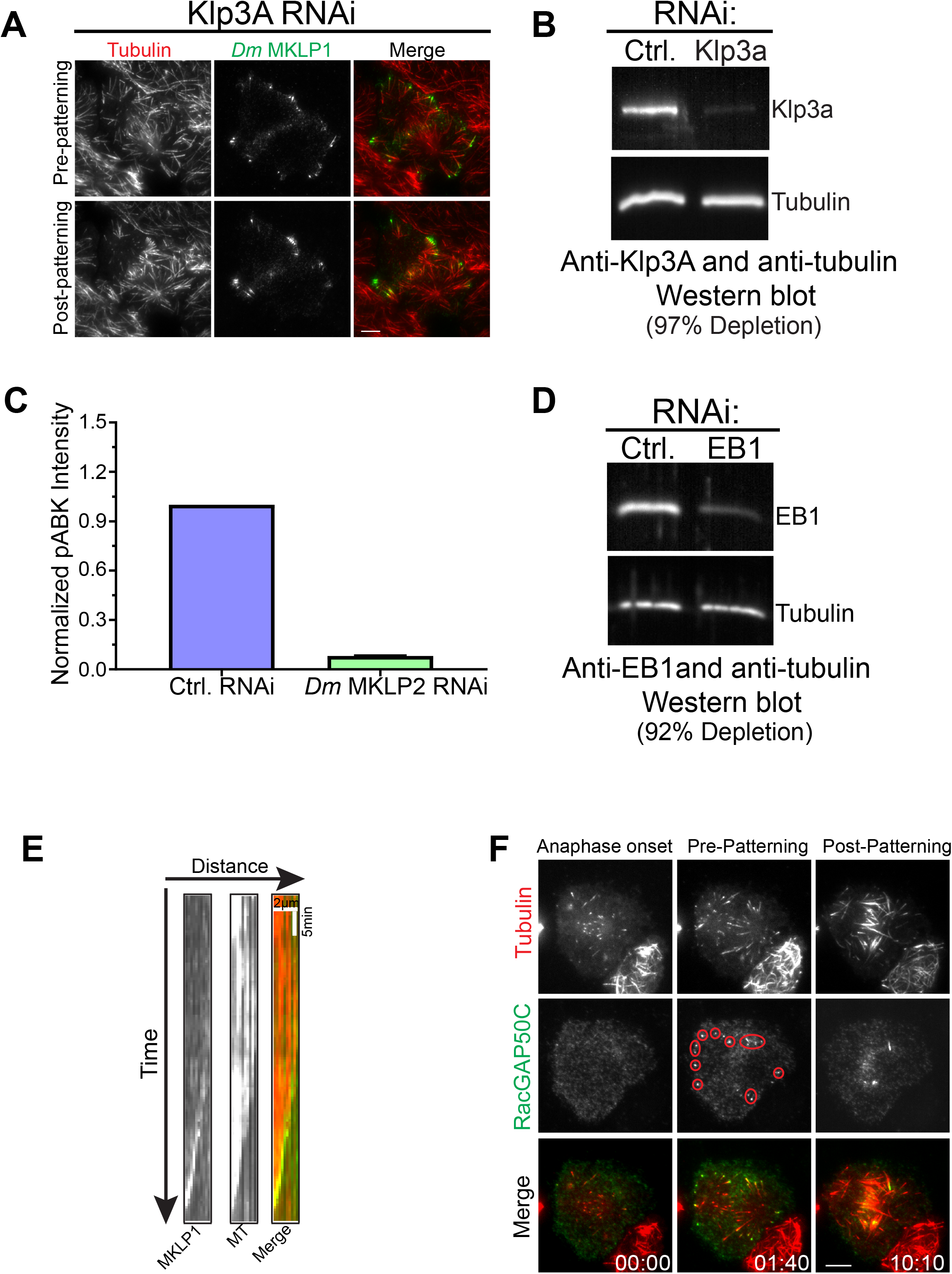
(A) Still-images from live-cell TIRF microscopy showing that *Dm* Kinesin-4 (Klp3A) depletion doesn’t affect CS-TIPs or midzone localization of *Dm* MKLP1 (red). (B, D) Western blots showing depletion of *Dm* Kinesin-4 (Klp3A) and EB1 by RNAi in cells expressing Polo-EGFP and MKLP1-EGFP respectively. (C) Quantitation of pABK intensity in control and *Dm* MKLP2 depleted cells, n=25, Error bar: SEM. (E) Kymograph of MKLP1 tip tracking (left) and microtubule polymerization (middle) generated from time-lapse TIRF microscopy (Supplementary Movie 1). The last image (right) shows the merge picture of MKLP1 and microtubule, Scale bar, 2μm, time: 1.5 min. (F) Still-images from live-cell TIRF microscopy showing RacGAP50C-EGFP (green) localization to the CS-TIPs (red) in control RNAi cells. Red ovals/circles highlight RacGAP50C localization to the CS-TIPs. Time: mins:secs. Scale bar, 5 μm.

**Supplementary Figure 2.**
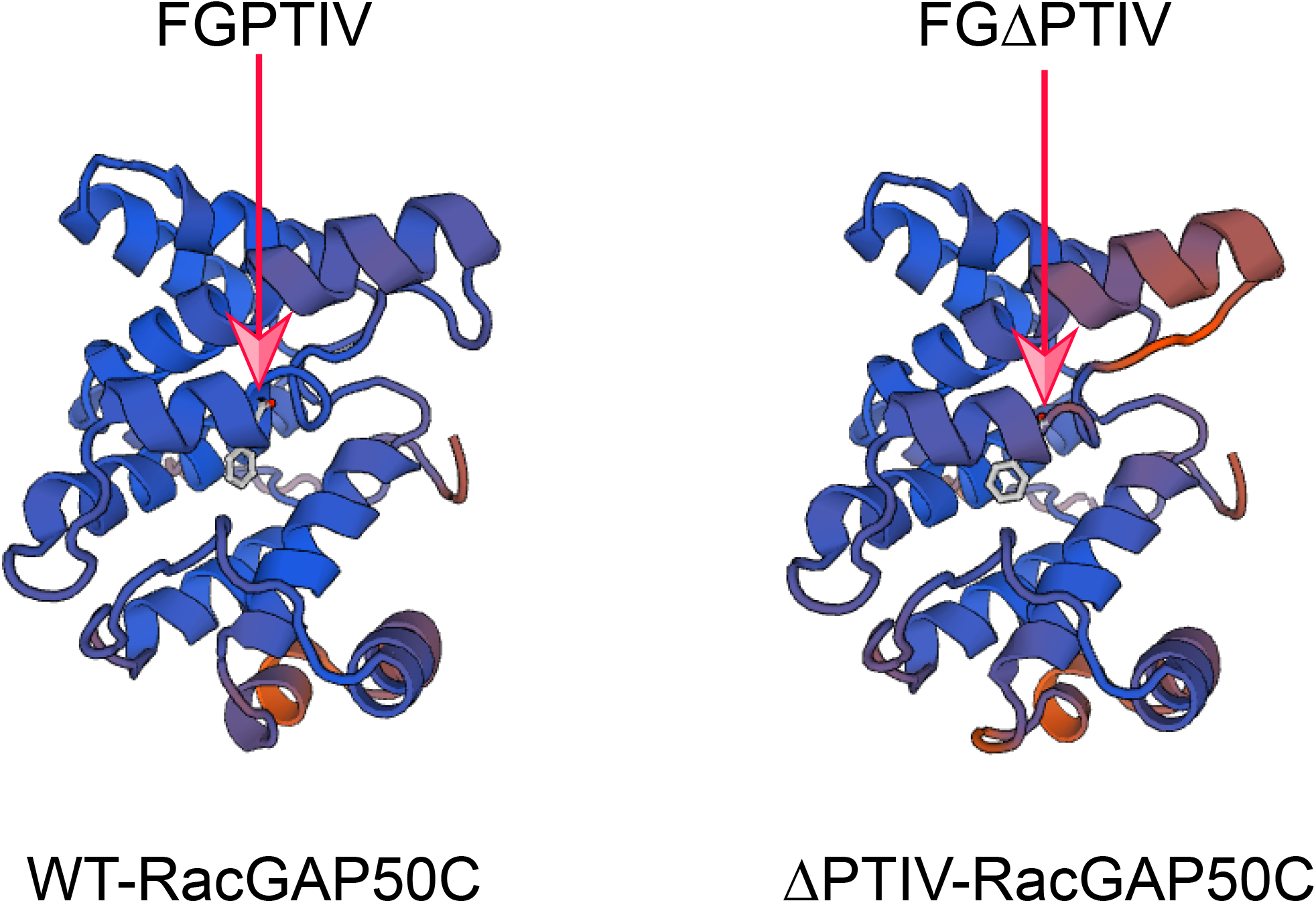
(A) Homology structure modelling of WT-RacGAP50C and ΔPTIV-RacGAP50C. Structures were generated using Rac GTPase-activating protein 1 (2ovj. 1. A) as a template in SWISS-MODEL. Arrow heads indicate the position of hxxPTxh motif (FGPTIV) in *Drosophila* RacGAP50C.

## SUPPLEMENTARY VIDEOS

**Video 1. *Dm* MKLP1 localizes to the CS-TIPs of polar and equatorial MTs.** Video shows co-expression of *Dm* MKLP1-EGFP (green) and Tag-RFP-α-tubulin (red) in S2 cells. *Dm* MKLP1-EGFP decorates the polar and equatorial CS-TIPs within ∼ 2-3 minutes of anaphase onset before becoming patterned onto equatorial MTs (∼7-10 minutes). *Dm* MKLP1-EGFP is also visible on the growing MT plus-tips and the central spindle. Frames were acquired at 10-sec. intervals on a Nikon Eclipse Ti-E - TIRF microscope. The playback rate is 10 frames per second. Time: mins:secs. Scale bar, 5 μm.

**Video 2. RacGAP50C localizes to the CS-TIPs of polar and equatorial MTs.** Video shows co-expression of RacGAP50C-EGFP (green) and Tag-RFP-α-tubulin (red) in S2 cells. RacGAP50C-EGFP decorates the CS-TIPs within ∼2 minutes of anaphase onset before becoming patterned onto equatorial MTs. RacGAP50C-EGFP is also visible on the growing MT plus-tips and the central spindle. Frames were acquired at 5-sec. intervals after anaphase onset on a Nikon Eclipse Ti-E - TIRF microscope. The playback rate is 10 frames per second Time: mins:secs. Scale bar, 5 μm.

**Video 3. Polo kinase localizes to the CS-TIPs and triggers cortical contractility.** Video shows the expression of Polo-EGFP in S2 cells. Polo-EGFP localizes to the CS-TIPs, central spindle, and interphase MTs. Cortical contractility and membrane invagination is clearly visible in the regions where CS-TIPs contact the cortex. Frames were acquired at 15-sec. intervals on a Nikon Eclipse Ti-E - TIRF microscope. The playback rate is 10 frames per second. Time: mins:secs. Scale bar, 5 μm.

**Video 4. *Dm* ECT2 (Pebble) accumulation on the CS-TIPs causes its auto-amplification near the cortex.** Video shows co-expression of *Dm* ECT2-EGFP (green) and Tag-RFP-α-tubulin (red) in *Drosophila* S2 cells. *Dm* ECT2 localization to the CS-TIPs at t = 3.30, and its amplification between t = 3.30 and 8.30 is clearly visible around the cortex. Accumulation of *Dm* ECT2-EGFP on midzone MTs and cortex is also visible. Frames were acquired at 5-sec intervals after anaphase onset on a Nikon Eclipse Ti-E - TIRF microscope. The playback rate is 20 frames per second. Time: mins:secs. Scale bar, 5 μm.

**Video 5. CS-TIPs activate RhoA.** Video shows co-expression of EGFP-Rhotekin (green) and Tag-RFP-α-tubulin (red) in *Drosophila* S2 cells. RhoA activation near the CS-TIPs is visible between t = 02:01 and 5:31. Cortical localization of Rhotekin-EGFP precedes its midzone localization. The midzone localization of Rhotekin-EGFP is visible between t = 5:31 and 07:31. Frames were acquired at 5-sec. intervals after anaphase onset on a Nikon Eclipse Ti-E - TIRF microscope. The playback rate is 10 frames per second.Time: mins:secs. Scale bar, 5 μm.

**Video 6. Polo-positive CS-TIPs activate RhoA.** Video shows co-expression of RFP-Rhotekin (red) and Polo-EGFP (green) in *Drosophila* S2 cells. In this movie, Polo-EGFP serves as a CS-TIPs marker. RhoA activation near the polo positive CS-TIPs is visible between t= 01:36 and 6:36. Frames were acquired at 5-sec. intervals after anaphase onset on a Nikon Eclipse Ti-E - TIRF microscope. The playback rate is 10 frames per second. Time: mins:secs. Scale bar, 5 μm.

**Video 7. CS-TIPs trigger cortical contractility by myosin accumulation.** Video shows co-expression of myosin light regulatory chain (MRLC)-RFP (red) and Polo-EGFP (green) in *Drosophila* S2 cells. Myosin accumulation and cortical contractility near the polo positive CS-TIPs are visible between t = 02:00 and 13:16. Frames were acquired at 5-sec. intervals on a Nikon Eclipse Ti-E - TIRF microscope. The playback rate is 10 frames per second. Time: mins:secs. Scale bar, 5 μm.

**Video 8. ABK is required for CS-TIPs assembly.** Video shows the expression of Polo-EGFP in S2 cells. Addition of ABK inhibitor, Binucleine 2, between t = 03:15 and 04:35 results in disassembly of CS-TIPs. Frames were acquired at 15-sec. intervals on a Nikon Eclipse Ti-E - TIRF microscope. The playback rate is 10 frames per second. Time: mins:secs. Scale bar, 5 μm.

**Video 9. ABK is required for CS-TIPs assembly.** Video shows the expression of *Dm* MKLP1-EGFP in S2 cells. Addition of ABK inhibitor, Binucleine 2, between t = 04:10 and 6:10 results in disassembly of CS-TIPs. Frames were acquired at 10-sec. intervals on a Nikon Eclipse Ti-E - TIRF microscope. The playback rate is 10 frames per second. Time: mins:secs. Scale bar, 5 μm.

**Video 10. Polo kinase is not required for CS-TIPs assembly.** Video shows the expression of Polo-EGFP in S2 cells. Addition of Polo inhibitor, BI 2536, between t = 02:45 and 03:53, doesn’t result in CS-TIPs disassembly. Frames were acquired at 15-sec. intervals after anaphase onset on a Nikon Eclipse Ti-E - TIRF microscope. The playback rate is 10 frames per second. Time: mins:secs. Scale bar, 5 μm.

**Video 11. Polo kinase is not required for CS-TIPs assembly.** Video shows co-expression of *Dm* MKLP1-EGFP (green) and Tag-RFP-α-tubulin (red) in S2 cells. Addition of Polo inhibitor, BI 2536, between t = 01:20 and 02:50, doesn’t result in CS-TIPs disassembly. Frames were acquired at 5-sec. intervals on a Nikon Eclipse Ti-E - TIRF microscope. The playback rate is 10 frames per second. Time: mins:secs. Scale bar, 5 μm.

**Video 12. Polo Kinase doesn’t localize to the midzone, but assembles normally on the CS-TIPs in Klp3A depleted cells.** Video shows the expression of Polo-EGFP in Klp3A depleted cells in *Drosophila* S2 cells. Polo-EGFP doesn’t localize to the midzone, but assembles normally on the CS-TIPs. Frames were acquired at 5-sec. intervals on a Nikon Eclipse Ti-E - TIRF microscope. The playback rate is 10 frames per second. Time: mins:secs. Scale bar, 5 μm.

**Video 13. ABK is required for CS-TIP signaling.** Video shows co-expression of EGFP-Rhotekin (green) and Tag-RFP-α-tubulin (red) in *Drosophila* S2 cells. RhoA activation near the CS-TIPs is visible between t = 02:00 to 3:55. Addition of ABK inhibitor, Binucleine 2, between t = 03:55 and 5:08 results in loss of Rhotekin from the CS-TIPs within ∼ 3 minutes. However, midzone Rhotekin signals take ∼ 6.30 minutes to disappear. Frames were acquired at 5-sec. intervals after anaphase onset on a Nikon Eclipse Ti-E - TIRF microscope. The playback rate is 10 frames per second. Time: mins:secs. Scale bar, 5 μm.

**Video 14. Polo kinase activity is required for CS-TIPs signaling.** Video shows co-expression of Rhotekin-EGFP (green) and Tag-RFP-α-tubulin (red) in *Drosophila* S2 cells. RhoA activation near the CS-TIPs is visible between t = 00:00 to 01:40. Addition of Polo inhibitor, BI 2536, between t = 01:40 and 2:51 results in loss of Rhotekin from the CS-TIPs within ∼ 3 minutes. Frames were acquired at 5-sec. intervals on a Nikon Eclipse Ti-E - TIRF microscope. The playback rate is 10 frames per second. Time: mins:secs. Scale bar, 5 μm.

**Video 15. ABK is required for CS-TIPs signaling.** Video shows the expression of MKLP1-EGFP (green) and Rhotekin-RFP (red) in *Drosophila* S2 cells. RhoA is visible around the cortex at t=1.00, but it gets reorganized near the CS-TIPs by t=3:58. Addition of ABK inhibitor, Binucleine 2, between t = 04:48 and 6:30 results in loss of Rhotekin from the CS-TIPs within ∼ 3 minutes. Frames were acquired at 5-sec. intervals after anaphase onset on a Nikon Eclipse Ti-E - TIRF microscope. The playback rate is 10 frames per second. Time: mins:secs. Scale bar, 5 μm.

**Video 16. Polo kinase activity is required for CS-TIPs signaling but not for assembly.** Video shows the expression of MKLP1-EGFP (green) and RFP-Rhotekin (red) in *Drosophila* S2 cells. RhoA accumulation near the CS-TIPs is visible after t = 01:02. Addition of Polo inhibitor, BI 2536, between t = 01:27 and 02:55 results in loss of Rhotekin from the CS-TIPs within ∼ 4.30 minutes, however MKLP1 localization to the cortex is still visible. Frames were acquired at 5- sec. intervals on a Nikon Eclipse Ti-E - TIRF microscope. The playback rate is 10 frames per second. Time: mins:secs. Scale bar, 5 μm.

**Video 17. Polo kinase activity is required for ECT2 accumulation near the cortex.** Video shows co-expression of *Dm* ECT2-EGFP (green) and Tag-RFP-T α-tubulin (red) in *Drosophila* S2 cells. ECT2 localization on the CS-TIPs is visible after t = 04:29. Addition of Polo inhibitor, BI 2536, between t = 04:34 and 05:49 results in loss of ECT2 from the CS-TIPs within ∼3 minutes. Frames were acquired at 5-sec. intervals after anaphase onset on a Nikon Eclipse Ti-E - TIRF microscope. The playback rate is 10 frames per second. Time: mins:secs. Scale bar, 5 μm.

**Video 18. EB1 is required for CS-TIPs assembly.** Video shows co-expression of *Dm* MKLP1-EGFP (green) and Tag-RFP-α-tubulin (red) in *Drosophila* S2 cells. In contrast to control RNAi cells, EB1 RNAi in cells expressing *Dm* MKLP1-EGFP results in loss of CS-TIPs assembly. Frames were acquired at 5-sec intervals on a Nikon Eclipse Ti-E - TIRF microscope. The playback rate is 10 frames per second. Time: mins:secs. Scale bar, 5 μm.

**Video 19. C-terminal truncation of RacGAP50C results in loss of CS-TIPs assembly.** Video shows co-expression of C-terminal truncation mutant of RacGAP50C-EGFP (green) and Tag-RFP-α-tubulin (red) in *Drosophila* S2 cells. In contrast to full length RacGAP50C, C-terminal truncation of RacGAP50C results in loss CS-TIPs assembly. Frames were acquired at 5-sec. intervals after anaphase onset on a Nikon Eclipse Ti-E - TIRF microscope. The playback rate is 10 frames per second. Time: mins:secs. Scale bar, 5 μm.

**Video 20. Deletion of ‘’PTIV” motif in RacGAP50C results in loss of CS-TIPs assembly.** Video shows co-expression of ‘’PTIV” deletion mutant of RacGAP50C-EGFP (green) and Tag-RFP-α-tubulin (red) in *Drosophila* S2 cells. In contrast to full length RacGAP50C, ‘’PTIV” deletion mutant of RacGAP50C results in loss CS-TIPs assembly. Frames were acquired at 5- sec. intervals after anaphase onset on a Nikon Eclipse Ti-E - TIRF microscope. The playback rate is 10 frames per second. Time: mins:secs. Scale bar, 5 μm.

